# Ultrastructural view of astrocyte-astrocyte and astrocyte-synapse contacts within the hippocampus

**DOI:** 10.1101/2020.10.28.358200

**Authors:** Conrad M. Kiyoshi, Sydney Aten, Emily P. Arzola, Jeremy A. Patterson, Anne T. Taylor, Yixing Du, Ally M. Guiher, Merna Philip, Elizabeth Gerviacio Camacho, Devin Mediratta, Kelsey Collins, Emily Benson, Grahame Kidd, David Terman, Min Zhou

## Abstract

Astrocytes branch out and make contact at their interfaces. However, the ultrastructural interactions of astrocytes and astrocytes with their surroundings, including the spatial-location selectivity of astrocyte-synapse contacts, remain unknown. Here, the branching architecture of three neighboring astrocytes, their contact interfaces, and their surrounding neurites and synapses have been traced and 3D reconstructed using serial block-face scanning electron microscopy (SBF-SEM). Our reconstructions reveal extensive reflexive, loop-like processes that serve as scaffolds to neurites and give rise to spongiform astrocytic morphology. At the astrocyte-astrocyte interface, a cluster of process-process contacts were identified, which biophysically explains the existence of low inter-astrocytic electrical resistance. Additionally, we found that synapses uniformly made contact with the entire astrocyte, from soma to terminal processes, and can be ensheathed by two neighboring astrocytes. Lastly, in contrast to densely packed vesicles at the synaptic boutons, vesicle-like structures were scant within astrocytes. Together, these ultrastructural details should expand our understanding of functional astrocyte-astrocyte and astrocyte-neuron interactions.

## Introduction

How astrocytes make contact with each other and with other constituents in the brain underlies the anatomic basis for astrocyte function in the central nervous system (CNS) (Barres, 2008; Clarke and Barres, 2013; Gomazkov, 2019). With respect to their anatomical organization, protoplasmic astrocytes are oriented in non-overlapping (i.e. distinct) domains (Bushong et al., 2002; Halassa et al., 2007; Ogata and Kosaka, 2002; Xu et al., 2014). Astrocytes within each domain can ensheath thousands of synapses within their occupancy volume using fine astrocytic processes (Peters et al., 1991; Wolff, 1970; Bushong et al., 2002; Khakh and Sofroniew, 2015). At the interfaces of astrocytic domains, gap junction coupling links astrocytes into a low resistance pathway in order to achieve syncytial isopotentiality, wherein neighboring astrocytes have the capacity to equalize their membrane potentials for high efficient regulation of brain homeostasis (Kiyoshi et al., 2018; Ma et al., 2016).

Given their ability to extensively contact neurites, astrocytes are integral components in the modulation of synaptic function (Papouin et al., 2017). Studies have shown that astrocytes are able to respond to synaptic events and regulate neuronal transmission (Adamsky et al., 2018; Henneberger et al., 2010; Jourdain et al., 2007; Panatier et al., 2011; Perea and Araque, 2007). Notably, astrocytes also extend their endfeet to make contact with blood vessels for uptake of nutrients needed for brain energy metabolism (Nortley and Attwell, 2017).

While these findings indicate the importance of astrocyte anatomy to function, much of the anatomic details at the ultrastructural levels remain to be resolved. For example, there is still much debate as to whether astrocyte branching architecture follows the classic ‘tree-like’ patterning (reminiscent of dendritic branches on neurons) or whether astrocyte processes form a reticular, sponge-like architectural design comprised of a meshwork of processes (Rusakov, 2015). Similarly, it is known that astrocytes only make contact at their domain interfaces with less than 10% of terminal processes (Bushong et al., 2002). However, an ultrastructural view of these inter-astrocyte terminal process contacts has yet to be resolved. Such information would provide critical anatomic insights into the low inter-astrocytic electrical resistance properties of astrocytes - a finding that was discovered by Kuffler and colleagues over half a century ago (Kuffler et al., 1966; Kuffler and Potter, 1964).

In addition to the unresolved questions regarding astrocyte branching patterns, astrocyte-astrocyte and astrocyte-synapse contacts also remain an important topic of investigation. To this end, it is not yet unknown whether synaptic support is solely carried out by the fine terminal processes, termed <u>p</u>erisynaptic <u>a</u>strocyte <u>p</u>rocesses (PAPs) (Peters et al., 1991; Wolff, 1970; Bushong et al., 2002; Khakh and Sofroniew, 2015), or whether every portion of the astrocyte is equally capable of providing structural synaptic coverage.

Critical to understanding these unanswered questions is the ability to resolve the ultrastructural complexity of astrocyte contacts. Indeed, fine astrocytic processes are structurally nanoscopic, which precludes the use of light microscopy and necessitates the use of electron microscopy (EM). In our current study, we’ve tackled these unanswered questions through a combined use of an *Aldh1l1*-eGFP transgenic mouse model and correlative confocal SBF-SEM techniques, wherein our EM specimen was confidently prepared to contain identity-validated and location-defined astrocytes, and preparation damage to the fine anatomic structures was significantly reduced - (Denk and Horstmann, 2004; Kiyoshi et al., 2018) - which is essential in order to resolve and reconstruct the fine astrocytic processes at nanoscopic ranges (Ventura and Harris, 1999).

In the current study, we examined - for the first time in requisite detail - the ultrastructure of an astrocyte connectome in an adult mouse hippocampus. The three reconstructed neighboring astrocytes allowed us to determine the branching pattern of astrocytic processes and identify inter-astrocytic contacts that altogether give rise to a low electrical resistance inter-astrocytic pathway. With the reconstruction of three across-astrocytic-domain neurites with their associated spines and contacting axons, we examined the spatial-location selectivity of astrocyte-synapse contacts, revealing a uniform distribution of synapses within and in-between astrocytic domains, and non-specific association of synapses with every physical part of an astrocyte. Taken together, our findings provide clarity to long-standing questions regarding the structural contacts within and between astrocytes, and they raise interesting functional questions regarding astrocyte-astrocyte and astrocyte-neurite interactions.

## Results

### Identification and 3D reconstruction of neighboring astrocytes using correlative confocal microscopy and SBF-SEM

To begin, we utilized *Aldh1l1*-eGFP reporter mice and confocal microscopy to define the region of interest (ROI) that contained three neighboring astrocytes in order to prepare the specimen for further SBF-SEM study. Specifically, after tissue fixation, low magnification images of coronal brain sections encompassing the cortex and hippocampus were taken. An angular cut made in the cortex serves as a point of reference for orientation purposes (Fig. 1A1). Note that we selected a tissue section that contained visible blood vessels which could be used as landmarks. An area containing eGFP+ astrocytes next to a visibly large blood vessel in the stratum radiatum of the hippocampus was chosen as our ROI (Fig. 1A2). The tissue section containing the ROI was then processed for SBF-SEM (Fig. 1B).

**Figure 1.**
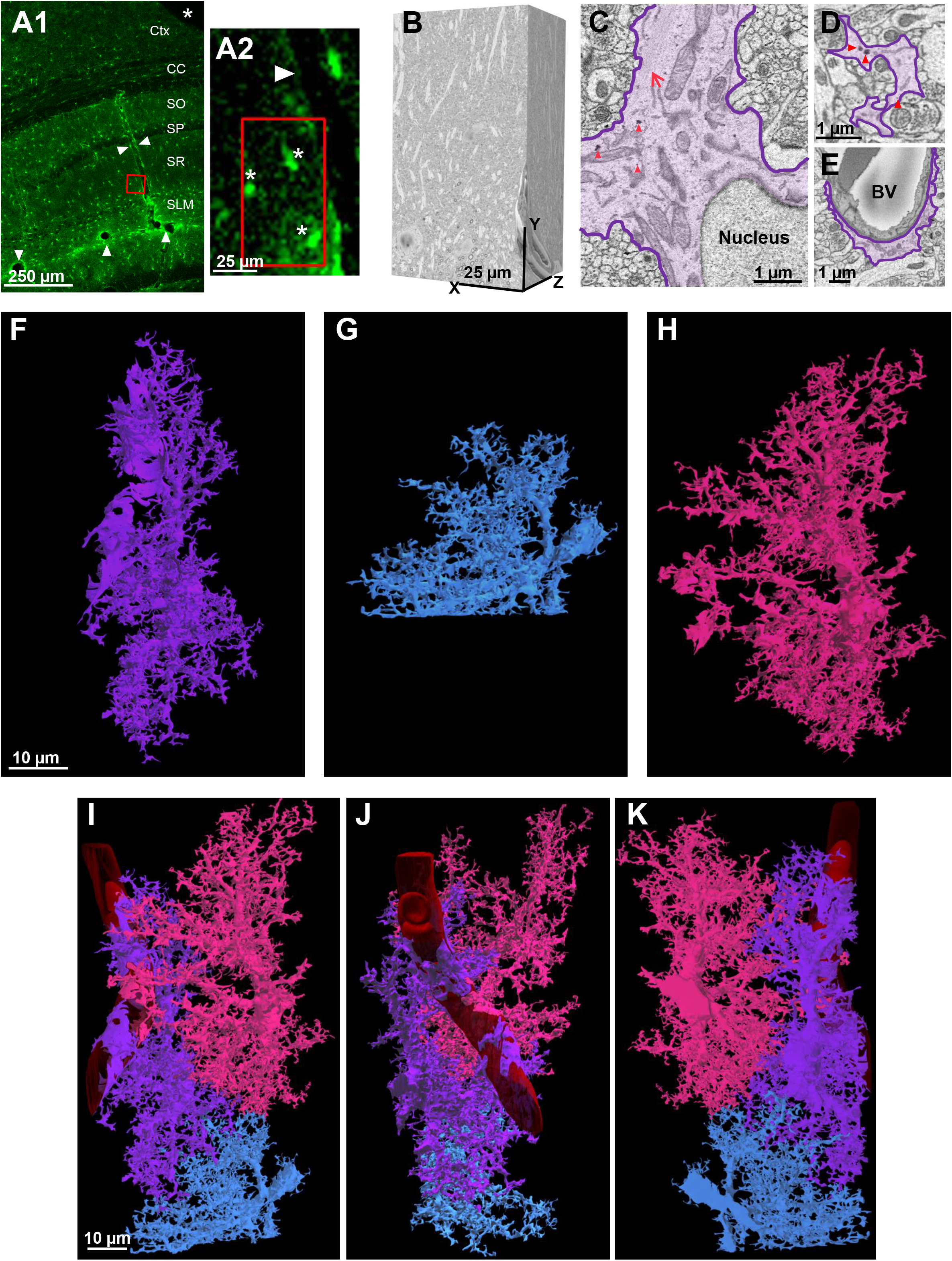
EM Identification and 3D reconstruction of neighboring astrocytes. **A1)** Low magnification confocal microscope image of a fixed brain section displaying the location of eGFP+ astrocytes and delineation of blood vessels (white arrowheads). An angular cut (*) at the upper-right region of the tissue serves as a fiduciary mark. **A2)** A magnified area from **A1** shows the spatial location of 3 neighboring astrocytes next to a blood vessel. Asterisks (*) denote the 3 astrocytes observed in the SBF-SEM images. **B)** The resulting 500-stack SBF-SEM volume dataset from the selected ROI containing neighboring astrocytes from the *stratum radiatum* hippocampal subregion. **C)** An astrocyte was identified by first locating the nucleus-containing cell body. Bundles of filaments (long, red arrow) and several examples of glycogen granules (red arrowhead, also see Fig. S1) are noted near the astrocytic nucleus. **(D)** Astrocyte processes (purple) that extend from the cell body possess an irregular and angular shape. **(E)** Processes that contact blood vessels expand into specialized astrocyte endfeet processes. Abbreviations: Ctx-Cortex, CC-Corpus callosum, SO-Stratum oriens, SP-Stratum pyramidale, SR-*stratum radiatum*, SLM-*stratum lacunosum-moleculare*, BV-blood vessel. **F-H)** 3-dimenstional view of three reconstructed astrocytes: purple, blue, and pink. **I-K)** Combined reconstruction depicting the front, side, and back views of the three astrocytes. Each astrocyte is labeled in a different color to clearly demarcate individual astrocyte domains and cellular structures. Note that the blue astrocyte appears ‘smaller’ in size as only part of the cell was included in the EM stack.

To identify astrocytes within the SBF-SEM dataset, we first located structures that were irregular and angular and shape and that showed visible cell bodies containing the nucleus (see Fig. 1C). We then examined the datasets to look for key structural characteristics of astrocytes, which included glycogen granules and bundles of intermediate filaments (Fig. 1C and D; see Fig. S1 for glycogen granule identification criteria). In addition, we followed the processes extending from the cell body, which formed well-defined astrocytic endfeet that make contact with blood vessels (Fig. 1E). Notably, these processes never made any distinct synaptic-like structures, therefore eliminating the possibility that they were neurons or NG2 glia (Bergles et al., 2000).

Within the SBF-SEM dataset, we identified three astrocytes (colored purple, blue, pink) and traced these astrocytes (and a surrounding blood vessel) to completion. 3D reconstructions were then created (Fig. 1F-H). Of note, the bottom half of the blue astrocyte was cut-off in the data set (Fig. 1G) and is therefore smaller in appearance. Overlaying the three astrocytes created a 3D reconstruction of an astrocyte connectome (Fig. I-K; Video S1).

### Characterization of astrocyte branching architecture

Previous reports utilizing light and electron microscopy have revealed that protoplasmic astrocytes possess a meshwork of small processes (Bushong et al., 2002; Shigetomi et al., 2013; Ventura and Harris, 1999); however, the intricate cellular architecture of astrocytes remains poorly understood. To examine this question, we used our astrocytes (reconstructed from our SBF-SEM dataset) to observe nanoscopic characteristics of these branching patterns (Fig. 2). In determining the branching architecture, we adopted the Root-Intermediate-Terminal (RIT) process labeling scheme, which was first used for dendritic arborization analysis (Uylings and van Pelt, 2002). This topography most closely resembled what we observed in our 2D astrocyte traces (Fig. 2A1-C1) and in our 3D reconstructions (Fig. 2A2-C2). Using this organizational scheme, we defined a *root* process as a branch that originates from the soma (Fig. 2A1-A2). *Intermediate* processes are those that branch from the root processes and on certain occasions, extend to other intermediate processes (Fig. 2B1-B2). Finally, small, thin *terminal* processes extended from the soma, root processes, and intermediate processes (Fig. 2C1-C2). These terminal processes did not progress any further and helped to delineate the astrocyte domain border.

**Figure 2.**
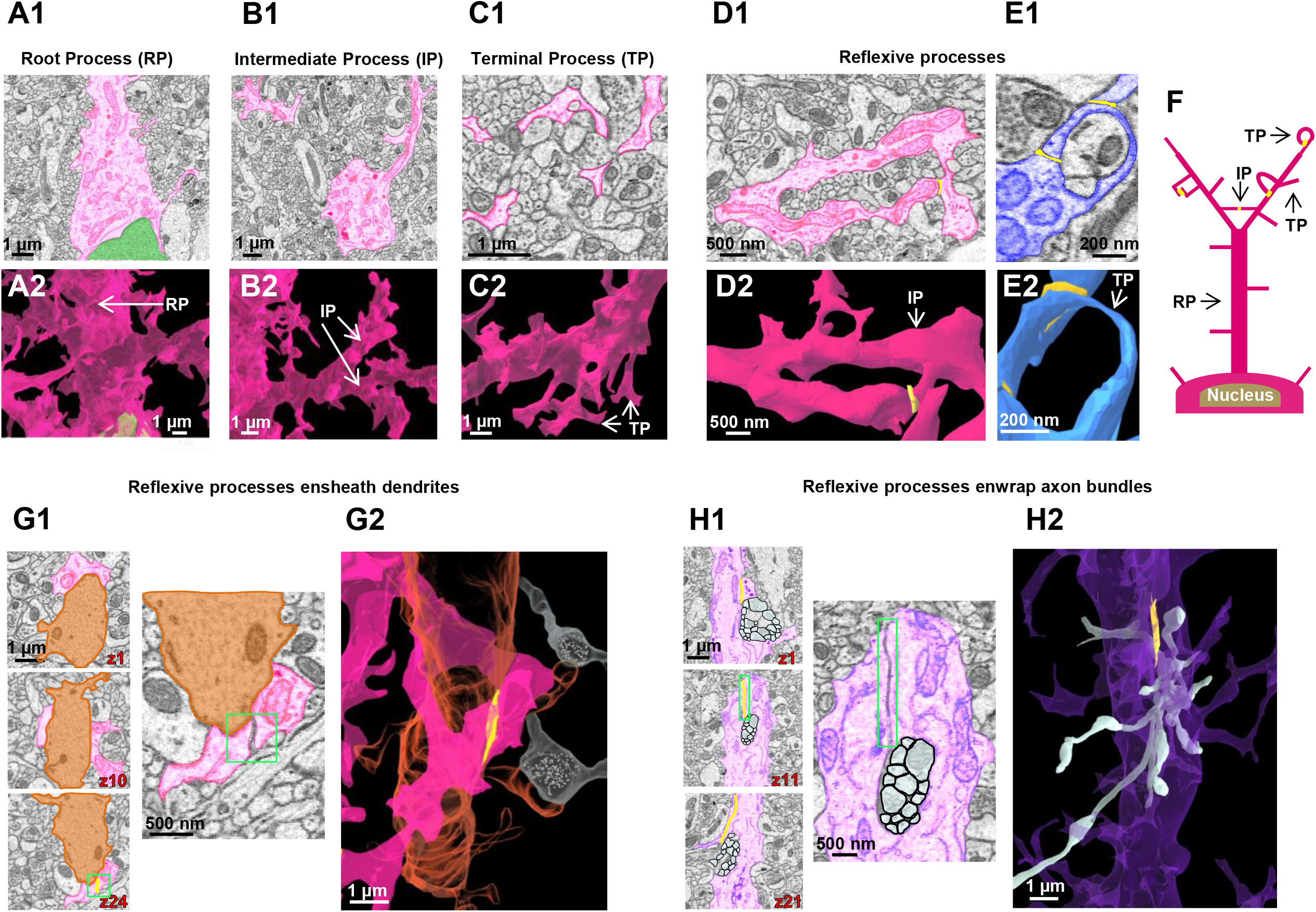
Astrocyte processes and their structural support to neurites. **A1)** 2D EM trace of an astrocyte root process. Root processes stem directly from the cell body of the astrocyte. The astrocyte nucleus is labeled in green. **A2)** Reconstructed 3D view of an astrocyte root extending from the cell body. Similar to images in the top panels, the nucleus is shown in green. **B1)** 2D EM trace of an astrocyte intermediate process extending from the root process. **B2)** Reconstructed 3D view of astrocyte intermediate processes branching out from the root process. **C1)** 2D EM trace of terminal astrocyte processes. **C2)** Reconstructed 3D view of astrocyte terminal processes extending from an intermediate process. Note that terminal processes are smaller than root and intermediate process; they reach a terminal (endpoint) and do not branch any further. **D1)** 2D EM image and 3D reconstruction **(D2)** of an intermediate process leading to a blood vessel. This process branches into two intermediate processes, which creates a reflexive contact near the endfoot process. Reflexive contacts are indicated in yellow in both the 2D images in the 3D images. **E1)** An EM image and 3D reconstruction **(E2)** of a single terminal process looping back to the intermediate process. A distal terminal process makes a reflexive contact near the top of the loop. Note that these reflexive processes are not completely fused (i.e. they do not form closed ‘loop’ structures). **F)** Simplified schematic diagram depicting the astrocyte branching architecture and reflexive processes. Contacts are noted in yellow. Abbreviations - IP: intermediate process; TP: terminal process; RP: root process. **G1)** Serial 2D EM images of astrocyte intermediate processes (pink) encircling a dendritic shaft (orange). The bottom image in the left panel exhibits a reflexive contact (boxed in green). Higher magnification images in the right panel clearly show the contact depicted in z slice 24. **G2)** 3D reconstruction showing the astrocyte intermediate process (pink) wrap around a dendrite (orange). Two axons (white) in synaptic contact with the associated dendritic spines are also represented. **H1)** 2D serial EM images of a branch point at the end of an astrocyte root process (purple) that splits into two intermediate processes, which enwrap a bundle of axons (white). Note the astrocyte reflexive contact in the middle panel (magnified on right side). **H2)** Side view of a 3D reconstruction of the axons (white) protruding through the enwrapment of the purple astrocyte process. All 3D reconstructions throughout the figure originated from the same astrocyte regions depicted in the corresponding 2D EM traces.

### Nanoscopic depiction of reflexive astrocyte processes and their structural support to neurites

Neurons typically display dendritic architecture, exhibiting tree-like arborizations wherein different branches/processes do not self-connect or form loop-like structures. In contrast, astrocytes exhibit spongiform morphology which can consist of processes that form connections within the same cell (Giaume et al., 2010). However, it is unclear whether this dense meshwork of processes can form cytoplasmic, fused loop-like structures that then make direct self-connections (i.e. reflexive contacts)(Rusakov, 2015).

Our 2D SBF-SEM traces and 3D reconstructions revealed that astrocyte processes make reflexive contacts within the same cell (Fig. 2D-E). We found that intermediate processes could create reflexive contacts wherein two ‘daughter processes’ split at a branch point and then reconnect (Fig. 2D1-2). Further, thin terminal processes that extended from an intermediate process also created reflexive contacts by ‘looping’ around to contact the intermediate process from which it originated (Fig. 2E1-2). Reflexive processes were widespread within the spongiform astrocyte domain, and it is interesting to note that these reflexive processes resemble the ‘O-ring’ structures observed in super-resolution imaging studies of hippocampal astrocytes (Arizono et al., 2020; Panatier et al., 2014); however, these reflexive processes were not completely fused and exhibited well-defined membrane borders (see middle panels in Fig. 2G1 and H1 and the 3D reconstructions in Fig. 2G2-H2; also see yellow reflexive contacts in Videos S2 and S3).

Astrocyte processes are extensively interwoven with surrounding neurites. To gain insight into the ultrastructural relationship between both cell types, we traced and reconstructed astrocytes in association with neural processes. We observed an intermediate astrocyte process enwrap an entire dendrite, and then form a reflexive contact to allow the intermediate process to further extend (Fig. 2G1-G2; Video S2). Additionally, we observed a root process enwrap neurites, where it split in a similar manner and formed a reflexive contact around a bundle of axons (Fig. 2H1-H2; Video S3). We also observed several examples of terminal processes enwrapping neurites (such as the image shown in Fig. 2E). These observations favor the view that structural support is a key function of astrocytes.

### Ultrastructural visualization of inter-astrocyte contact patterns

Having characterized branching patterns *within* the same astrocyte, we next sought to examine branching/contact patterns *between* neighboring astrocytes. It is important to note that an astrocyte network is an electrically low-resistance pathway (Kuffler et al., 1966) that is critical in establishing syncytial isopotentiality, or the ability for coupled astrocytes to behave as a single unit (Kiyoshi et al., 2018; Ma et al., 2016). However, the anatomical characteristics of astrocytes that allow for this low-resistance functional feature have not yet been determined. Our 3D reconstructions of three neighboring astrocytes enabled us to map, for the first time, the connectivity patterns between astrocytes (Fig. 3A). To begin, we found that a process extending from one astrocyte could contact more than one neighboring astrocyte (Fig. S2). Additionally, we found that one astrocyte only made minimal overlap with a neighboring astrocyte - leaving a very narrow interface wherein the two astrocyte domains made contact (see yellow contacts in Fig. 3A). Furthermore, close examination of astrocytic terminal processes revealed multiple types of contacts between astrocytes (Fig. 3B-D). We classified these contacts into two distinct categories: point-point contacts (Fig. 3B; Video S4) and elongate contacts (Fig. 3C; Video S4). Point-point contacts displayed a small surface area between the ‘tips’ of the terminal processes. It is presumed that these point-point contacts are the site of gap junctions. Elongate contacts, on the other hand, showed a larger contact surface area.

**Figure 3.**
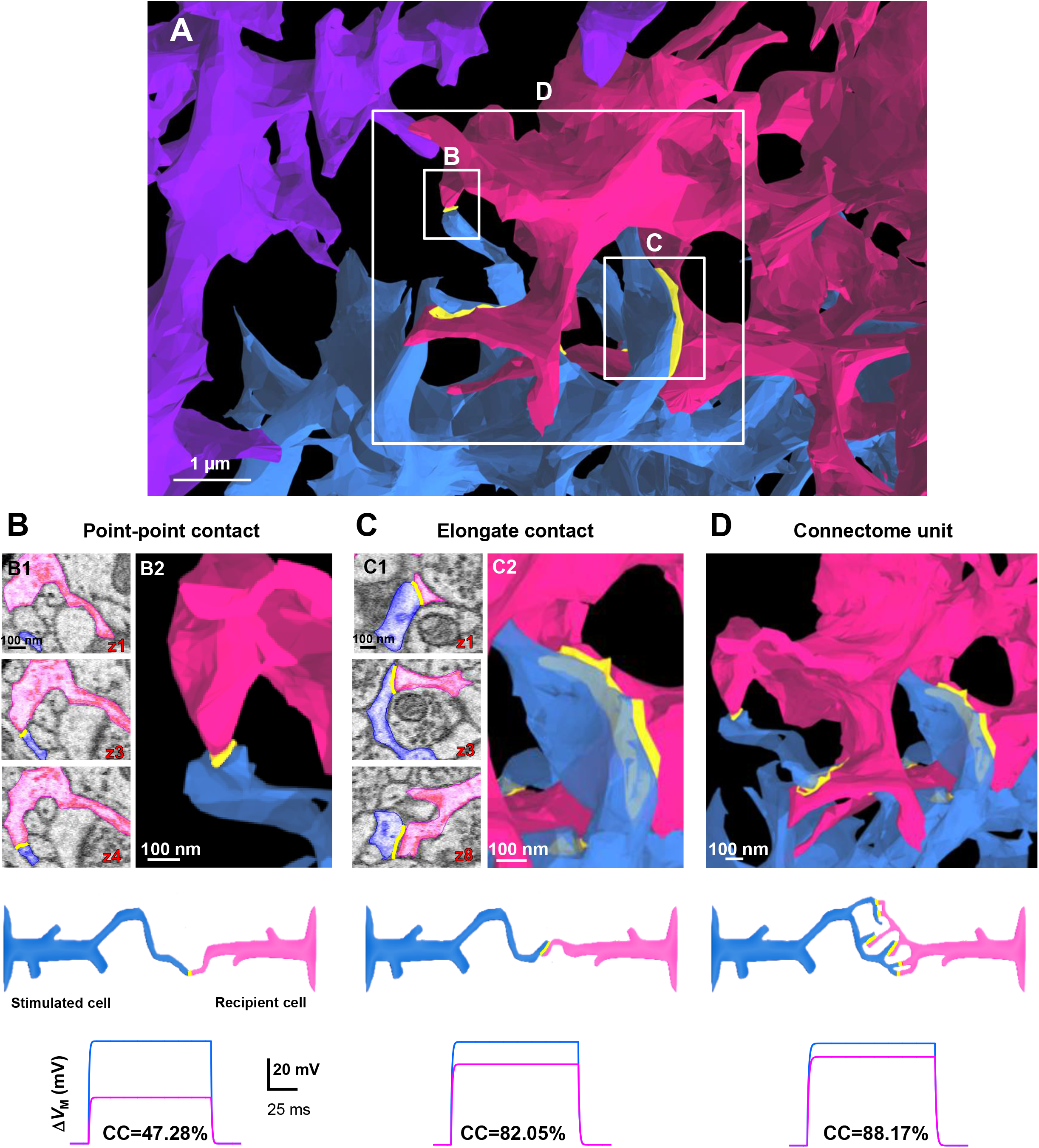
Contact patterns of astrocytes at their interfaces and computational modeling of pathway resistance in relation to astrocyte contact type. **(A)** 3D reconstruction of astrocyte processes reveals the interface between the astrocytes. For ease of viewing the ultrastructural inter-astrocyte contacts (between the pink and blue astrocytes), fewer intermediate and terminal processes were reconstructed in this image (relative to the images depicted in Fig. 1). Also note that contacts boxed in white are magnified in **B-D**. **B)** 2D serial EM images **(B1)** of point-point contacts between two astrocytes. These point-point contacts persist through a few serial sections to create a small contact at the tip of the terminal processes (see 3D EM image in **B2). C)** 2D serial EM images **(C1)** of an elongate contact. Elongate contacts typically contact/persist through many serial sections to create a large contact area (see 3D EM image in **C2)**. **D)** A 3D reconstruction of a ‘connectome unit’. This connectome unit contains a cluster of six mixed contacts (combination of point-point and elongate). Throughout the figure, note that all 2D traces are from the same astrocyte region used to generate the 3D reconstructed images. Schematic representations are provided at the bottom of the figure to illustrate single point-point contacts, elongate contacts, and a ‘connectome unit’. Further, the bottom panels represent coupling coefficient (CC) modeling. A single point-point contact yields a 47.28% CC **(B)**, whereas a single elongate contact **(C)** increases the CC to 82.05%. A full connectome unit consisting of a cluster of 2 elongate and 4 point-point contacts increased the CC to 88.17% **(D).**

Turning back to the idea of how low-resistance pathways are established, we found that many contact sites consisted of clusters of both point-point and elongate contacts. In one particular location, we observed astrocytes make six different contacts, and we termed this complex contact region as an astrocyte ‘connectome unit’ (Fig. 3D; Video S4). Within this connectome unit, multiple terminal processes extended from intermediate processes in a fork-like manner (Fig. 3D). We hypothesized that the multiple-contact anatomical characteristics of this ‘connectome unit’ serve as the inter-astrocytic electrical connectors that reduce the electrical resistance between astrocytes.

### Computational modeling of the astrocyte connectome simulates a lowered pathway resistance

In our recently published report (Ma et al., 2016), astrocyte syncytial isopotentiality was computationally modeled to estimate the strength of electrical coupling among various astrocytes. In this model, each astrocyte in the syncytium is displayed as a single compartment, and the requisite experimental conditions (i.e. ion concentrations and membrane permeabilities) were recreated. This model demonstrated the dependence of the coupling coefficient (CC) on the strength of gap junction coupling, wherein CC is defined as the ratio of the current-induced membrane potential (*V*_M_) of the recipient cell to the *V*_M_ of the stimulated cell (Ma et al., 2016). We expanded upon this model to create a spatial, multi-compartment representation that incorporates the ultrastructural anatomy of the reconstructed astrocyte connectome in relation to pathway resistance.

The CC in a pair of coupled astrocytes was determined to be 94% (Ma et al., 2016). We computationally modeled astrocyte contacts from Fig. 3 to examine the CC. The anatomical measurements of the processes leading to the contact interface and the process contact areas were incorporated into this spatial model. Further, the area between the processes was measured as the presumed site of gap junctions, and the number of gap junctions was estimated based on the density of gap junctions from freeze-fracture replica immunogold labeling experiments (Rash et al., 2001). A current injection of 0.5 nA for 150 ms was applied to the stimulated cell, and the *V*_M_ responses from the stimulated and recipient cells were recorded.

In our model, we found that a pathway consisting of only a single point-point contact constitutes a CC of 47.28% (Fig. 3B; bottom panel). A single elongate contact yields a larger CC at 82.05%, which is attributed to an increased number of gap junctions (Fig. 3C; bottom panel). Finally, the full cluster of six contacts (i.e. a ‘connectome unit’) increased the CC to 88.17% (Fig. 3D; bottom panel). In addition, our computational model further predicts that 2, 3, and 4 connectome units can further increase the CC values to 92.5%, 93.92%, and 94.35%, respectively (schematic data not shown). Hence, our model indicates that the use of multiple contacts/contact types is crucial in the ability of astrocytes to reduce electrical resistance.

### 3D reconstruction of astrocytes in association with neurites

Astrocyte processes are extensively interwoven with surrounding neurites. Reconstruction of three astrocytes in association with three neurites provided us with ultrastructural details of the anatomical relationships between both cell types (Fig. 4). We first traced (to completion) three dendrites and their associated spines and contacting axons (Fig. 4C; Video S5). To more easily view the dendritic spines, a small number of axons are depicted in Fig. S3A. Note the detailed structure of the dendritic spines shown in our reconstructions-representative of all six standard spine categories: thin, mushroom, stubby, cup, branched, and filopodia-like (Fig. S3B1-B6) (Hering and Sheng, 2001; Risher et al., 2014). The three neurite reconstructions were then combined with the 3D reconstructions of the three astrocytes in order to generate a complete network-level view of the astrocyte-neurite interaction (Fig. 4D; Video S6).

**Figure 4.**
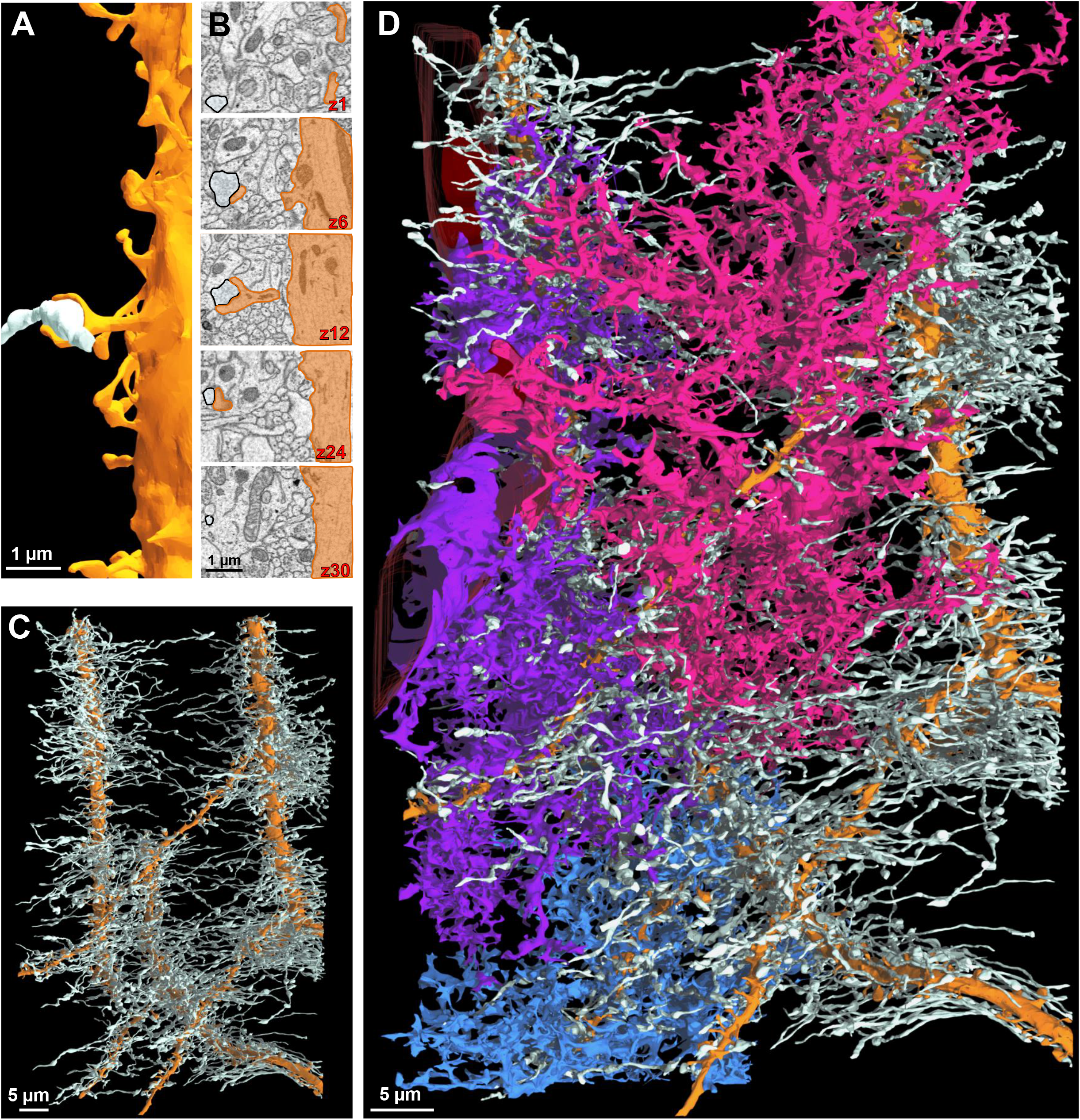
An ultrastructural view of astrocyte-neurite association. **A)** Partial 3D reconstruction of one dendrite. An axon (white) is drawn for reference in order to depict the axon-dendritic spine interface (i.e. synapse). **B)** 2D serial traces of the axon (white) and dendritic spine (orange) that form a synapse (synapse is depicted in **A** and in z-section 12). **C)** 3D reconstruction of three dendrites (orange) and their associated axons (white) shown in a front view. **D)** Front view of the three neighboring astrocytes and their association(s) with the three reconstructed neurites.

### Uniform association of synapses with astrocytes from soma to terminal processes

A current view is that astrocytes interact with synapses using their PAPs, or thin processes positioned next to synapses (Peters et al., 1991; Wolff, 1970). While the structural aspects of PAPs tend to be more characteristic of fine/terminal processes, we sought to examine whether the entire astrocyte domain could provide synaptic coverage. Indeed, we found that every physical part of an astrocyte can make contact with synapses (Fig. 5A). Ultrastructural 3D reconstructions of synaptic and pre-/post-synaptic elements that sat adjacent to the astrocyte soma (Fig. 5B1-2), root processes (Fig. 5C-2), intermediate processes (Fig. 5D1-2), terminal processes (Fig. 5E1-2), reflexive processes (Fig. 5F1-2), and even astrocytic endfeet (Fig. 5G1-2), provided clear evidence that all cellular compartments of the astrocyte provide synaptic coverage. Quantitative analysis of synapses that abut processes throughout the astrocyte domain revealed a significant effect of astrocyte process type on synapse density (Fig. 5H; F_(5 , 43)_ = 19.11; p < 0.0001; See Table S1 for post-hoc analysis), with the greatest density of synapses found to abut terminal and reflexive processes. Altogether, these synaptic associations demonstrate that all astrocyte compartments are capable of providing synaptic coverage.

**Figure 5:**
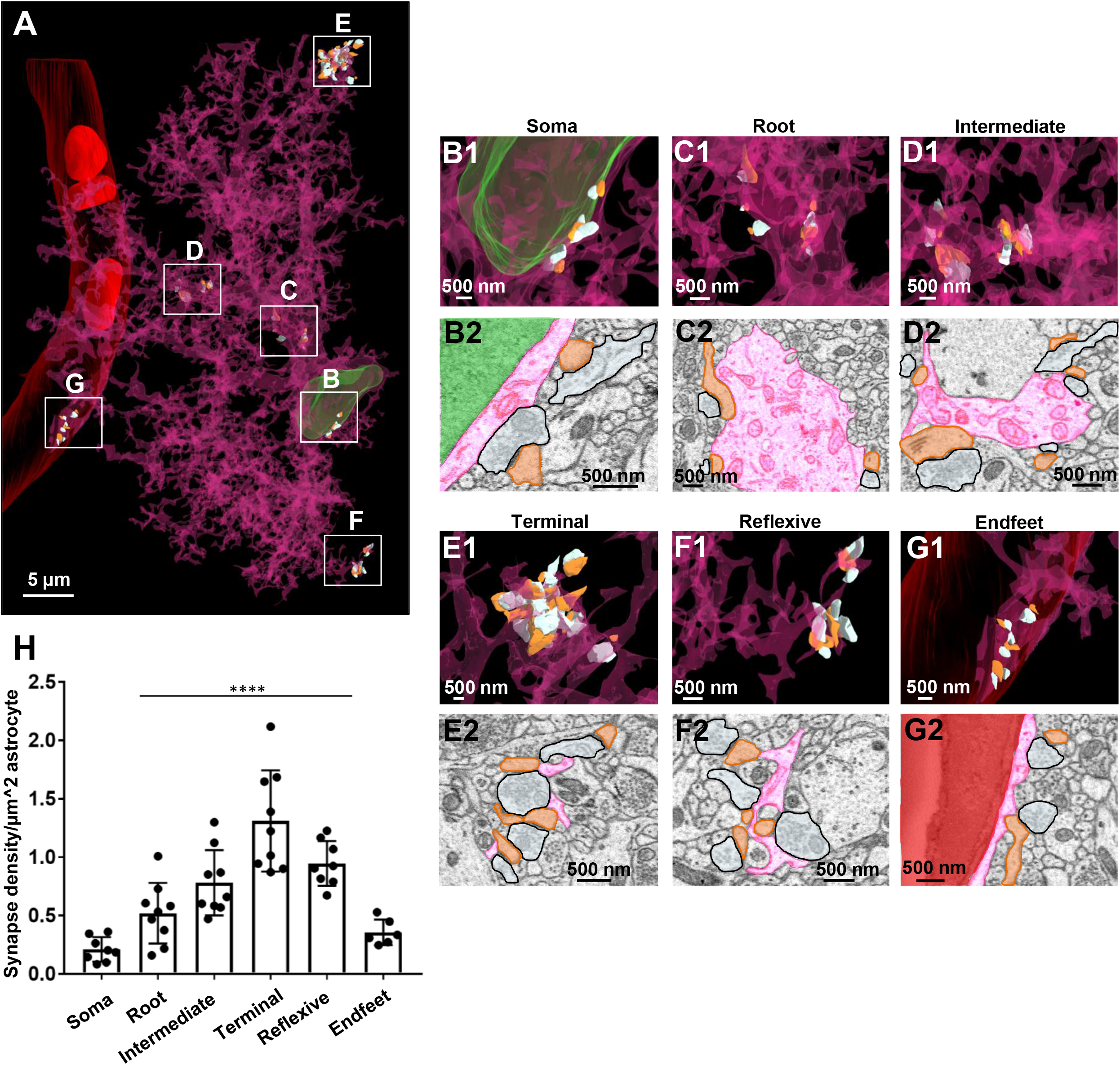
Enrichment of synapses abutting astrocytic terminal and reflexive processes. **A)** 3D reconstruction of an entire astrocyte (pink) and its contacts with synapses (orange: postsynapse and white: presynapse). Note that the white, boxed areas in ‘A’ approximate the locations of magnified images in **B-G**. Further, the synapses depicted in the representative image **(A)** for each region were constructed from approximately the same volume of astrocyte. **B1)** Magnified 3D reconstruction of multiple synapses contacting the astrocyte soma. **B2)** 2D EM trace of synapses contacting the astrocyte soma. Note that the astrocyte nucleus is depicted in green in both the 2D and 3D images. **C1)** 3D reconstruction and 2D trace **(C2)** of synapses contacting the root process of an astrocyte. **D1)** 3D reconstruction and 2D trace **(D2)** of synapses contacting intermediate processes of an astrocyte. **E1)** 3D reconstruction and 2D trace **(E2)** of synapses contacting terminal astrocytic processes. **F1)** 3D reconstruction and 2D trace **(F2)** of synapses contacting reflexive astrocyte processes. **G1)** 3D reconstruction and 2D trace **(G2)** of synapses contacting the astrocyte endfeet adjacent to the blood vessel (red). **H)** Graphical representation of the synapse density per area of astrocyte. Data was obtained from three reconstructed astrocytes, with three ROIs chosen per astrocyte process region (soma, root, intermediate, terminal, reflexive, endfeet); represented as mean + SEM. Note that one of three astrocytes (blue) did not contact a blood vessel, and thus only six ROIs are shown in the ‘endfoot’ panel of the graph. Also note that both intermediate processes and terminal processes that extend from intermediate processes were included in the ‘reflexive’ analysis. ****: p < 0.0001; One-way ANOVA, followed by post-hoc tests.

### The majority of synapses are ensheathed by astrocytic processes

Having established that all parts of an astrocyte domain can make contact with synapses (Fig. 5), we next examined their ultrastructural contacts. Several bodies of work examining both the developing and mature rat cortex and hippocampus have reported that a large number of synapses make contact with astrocyte processes (Kikuchi et al., 2020; Ventura and Harris, 1999; Witcher et al., 2007). However, to date, the percentage of synapses contacted by astrocyte processes within the mature mouse hippocampus is not known. Using the three fully reconstructed neurites (presented in Fig. 4), we analyzed the percentage of astrocytic processes at the axon-spine interface (i.e. synapse). In total, we evaluated 920 synapses from the three reconstructed neurites, and recorded whether astrocyte processes made contact with the synaptic cleft (Fig. 6A1-A2), pre or post synaptic elements (Fig. 6B1-B2), or if no astrocytes made contact with the synapse (Fig. 6C1-C2). Additionally, each synapse was classified as ‘asymmetric’ or ‘symmetric’ based on whether it exhibited a prominent or narrow postsynaptic density (Gray, 1959; Kikuchi et al., 2020; Peters and Palay, 1996; and see Fig. 6D1).

**Figure 6.**
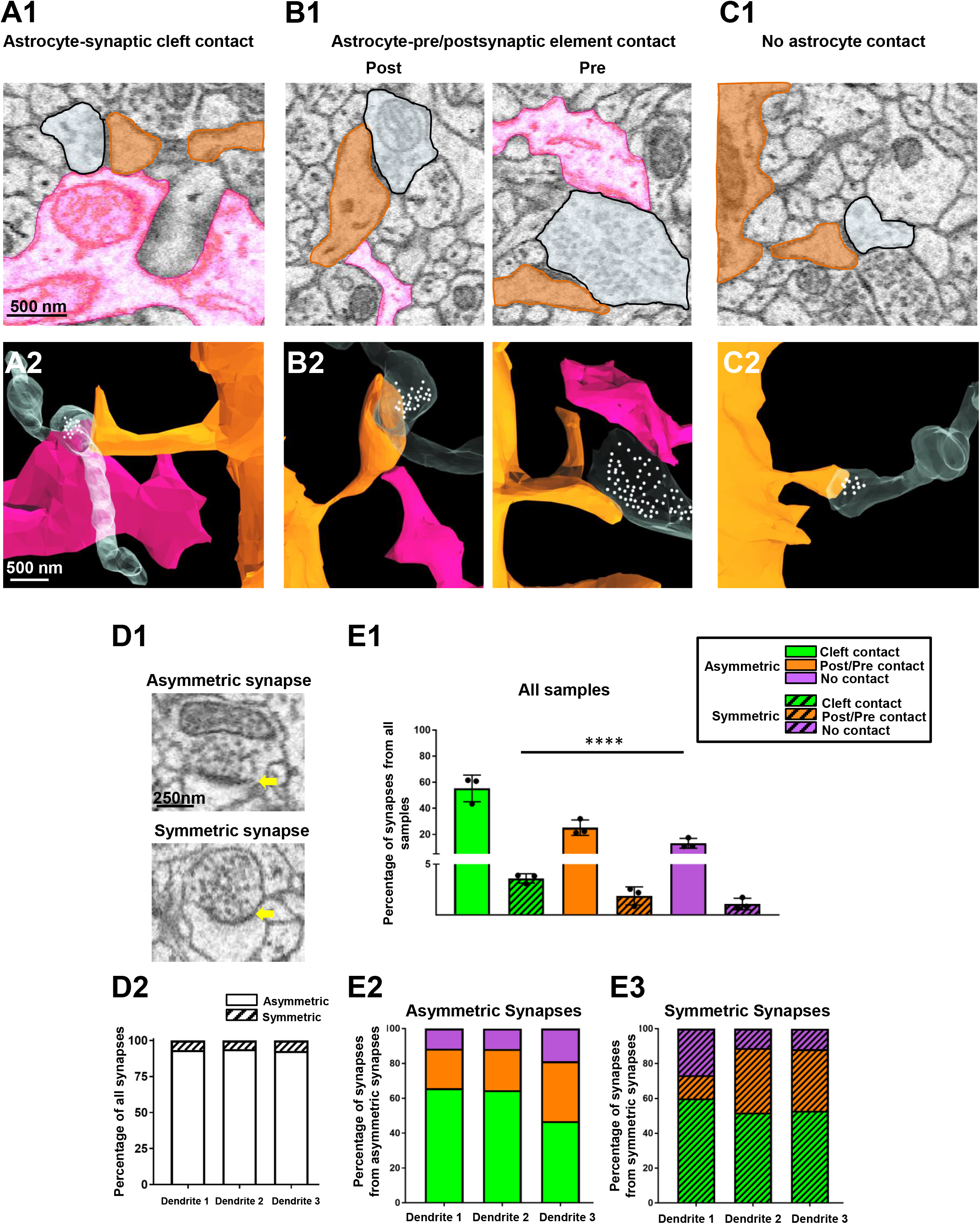
The majority of synapses are contacted by astrocytic processes. **A1)** 2D EM trace of an astrocyte process (pink) contacting the synaptic cleft. **A2)** 3D reconstruction from **A1. B1)** 2D EM traces of astrocyte process contacting either post-synaptic dendritic elements (left panel) or pre-synaptic elements (right panel). **B2)** 3D reconstruction from **B1**. **C1)** 2D EM trace of a synapse with no astrocyte contact. A 3D reconstruction is also depicted in **C2**. In all representative images, the astrocyte processes that contact the synapses are from one astrocyte (pink) and the synapses are from two fully reconstructed dendrites shown in Fig. 4. White spheres depict the approximate locations of synaptic vesicles observed from several serial 2D EM stacks. **D1)** 2D EM traces depict an example of an asymmetric synapse (prominent post synaptic density—top panel) and a symmetric synapse (modest post synaptic density— bottom panel). Yellow arrows denote the post-synaptic density. **D2)** Graphical representation of the percentage of asymmetric versus symmetric synapses (irrespective of astrocyte contact type) from all three traced dendrites. **E1)** Graphical representation of the percentage of synapses (asymmetric or symmetric) that contact astrocyte processes at the synaptic cleft, on pre or post synaptic elements, or have no contact with astrocyte processes. Note that these data were pooled from the synapses that contacted spines from all three reconstructed dendrites, and thus, each data point is representative of the percent coverage per dendrite. Data was analyzed by one-way ANOVA followed by post hoc tests; ****: p < 0.0001. **E2)** Distribution of the percentage of asymmetric synapses that have astrocyte contact with the synaptic cleft, post, or pre synaptic elements, or no astrocyte contact. **E3)** Distribution of the percentage of symmetric synapses that have astrocyte contact with the synaptic cleft, post or pre synaptic elements, or no astrocyte contact.

Assessment of asymmetric versus symmetric synapse percentage (irrespective of astrocyte contact type) revealed that a majority of synapses were asymmetric (Fig. 6D2). Along these lines, 218/235 synapses (93%) from dendrite 1, 216/231 synapses (94%) from dendrite 2, and 427/454 synapses (94%) from dendrite 3 were asymmetric. When examining synapse type (i.e. asymmetric and symmetric) with respect to astrocyte contacts, we found that a majority of synapses were contacted by astrocytes, with significant differences observed in astrocyte contact patterning (Fig. 6E1; F_(5, 12)_ = 51.08; p < 0.0001; one-way ANOVA. See Table S2 for post-hoc analysis). Specifically, 55% of all synapses were asymmetric and contacted astrocytes at the cleft (Fig. 6E1; solid green bar) and 4% of all synapses were symmetric and contacted astrocytes at the synaptic cleft (Fig. 6E1; patterned green bar). Further, 25% of all synapses were asymmetric and had pre- or post-synaptic contact with astrocytes (Fig. 6E1; solid orange bar) and 2% of all synapses were symmetric and had pre-or post-synaptic contact with astrocytes (Fig. 6E1; patterned orange bar). In contrast, 13% of all synapses were asymmetric and had no contact with astrocytes (Fig. 6E1; solid purple bar) and 1% of all synapses were symmetric and had no contact with astrocytes (Fig. 6E1; patterned purple bar). Individual dendrite-parsed graphical representations of asymmetric synapse-astrocyte contacts (Fig. 6E2) and symmetric synapse-astrocyte contacts (Fig. 6E3) are also shown. Taken together, these results suggest that most synapses (86% in total) have contact with astrocyte processes, and a majority of these astrocyte-synapse contacts occur at the synaptic cleft (compared to pre and/or post synaptic elements).

### Paucity of vesicles in astrocyte processes adjacent to synapses

Our extensive tracing of synapse-astrocyte contacts provided the opportunity for examination of intracellular vesicle-like structures that have been proposed to mediate gliotransmission-the transfer of information from glia to neurons (Araque et al., 1999; Perea et al., 2009).

To begin, we defined ‘vesicle-like organelles’ using synaptic vesicles as a reference (see Fig. 7A; for a description of criteria used to define vesicles, see Fig. S1 and STAR methods). With these criteria in mind, we examined all compartments of the astrocyte domain (Fig. 7B-F), given that each process type exhibited direct association with synapses (Fig. 5). Across astrocyte process types, we observed a paucity of vesicles positioned adjacent to synapses, in contrast to the massive assembly of vesicles noted in the presynaptic terminals (Fig. 7A-F). Though we did not observe an appreciable collection of vesicles in astrocyte processes (next to synapses), occasionally, we did observe a very sparse number of vesicle-like structures (see arrowheads in Fig. 7B, C, and E). Quantitative analysis of the density of vesicles per volume cubed of astrocyte (or neurite) confirmed our qualitative observations: indeed, in contrast to synaptic vesicles (see quantification of synaptic vesicle number-serving as a control in Fig. 7G), few astrocytic vesicles were observed across all compartments of the domain, with no significant difference in vesicle number between astrocyte process types (Fig. 7G; F_(4, 39)_ = 0.816; p = 0.523; one-way ANOVA). Together, the lack of apparent vesicles in astrocytes suggests that if gliotransmission occurs, it is likely not mediated through a Ca^2+^-dependent vesicular-release mechanism.

**Figure 7:**
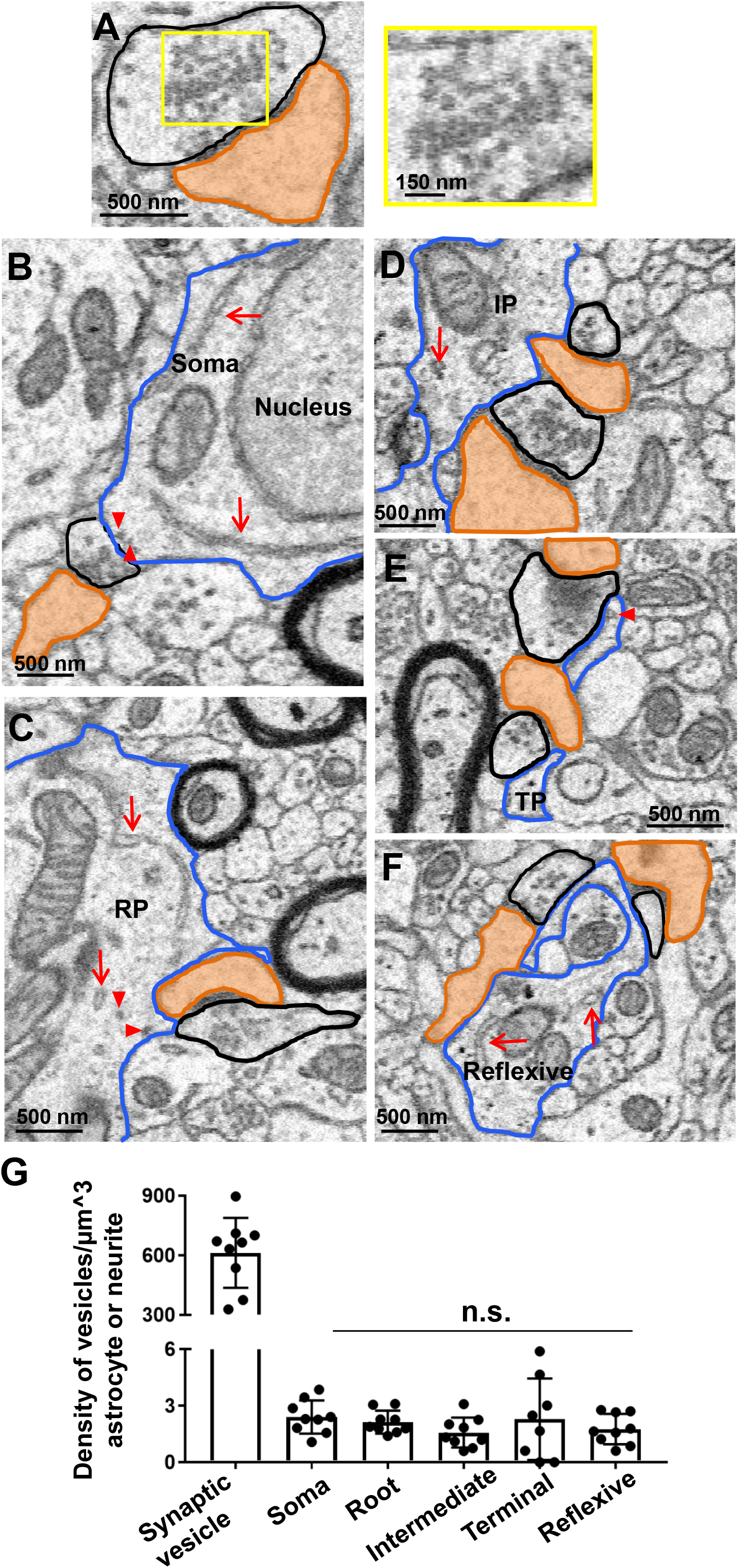
Paucity of vesicle-like structures in synapse-ensheathing astrocyte processes. **A)** Representative 2D EM image of a synapse (white: pre-synaptic element; orange: dendritic spine head). Note the significant number of vesicles in the pre-synaptic element (magnified in the right panel). **B)** 2D EM traces of an astrocyte soma (blue outline), **(C)** astrocyte root process (blue outline), **(D)** astrocyte intermediate process (blue outline), **(E)** astrocyte terminal process (blue outline), and **(F)** astrocyte reflexive process (blue outline). Synapses are also outlined (black: presynaptic element; orange: dendritic spine head). Note that very few (if any) vesicle-like structures were found in astrocyte processes that contacted synapses. Likely astrocyte vesicles are denoted by red arrowheads, and endoplasmic reticulum is denoted by long red arrows. **G)** Quantification of the approximate number of vesicles in each astrocyte (or neurite). Data is represented as mean + SEM. No significant difference in the density of vesicle-like structures was found between process types. n.s.: not significant. Abbreviations— RP: root process; IP: intermediate process; TP: terminal process.

### Neighboring astrocytes can share coverage of a single synapse

The extensive astrocytic coverage of synapses that we observed led us to examine whether synapses could be ensheathed by processes stemming from different astrocyte domains, or whether domain exclusivity of synaptic coverage (Halassa et al., 2007) holds true. We have evidence of a synapse receiving synaptic coverage by two neighboring astrocytes (Fig. 8; Video S7). Note that on each side of the synapse, a different astrocyte process (pink or blue) is adjacent to the synaptic cleft (Fig. 8A-B; Video S7). This coverage persisted across a significant portion of the synaptic cleft, and while it continued across multiple z-planes, it simultaneously formed contacts with the other astrocyte process (Fig. 8A; Video S7). These 2D traces and 3D reconstructions suggest that the interaction of synapse with astrocytic process is not restricted within a domain.

**Figure 8:**
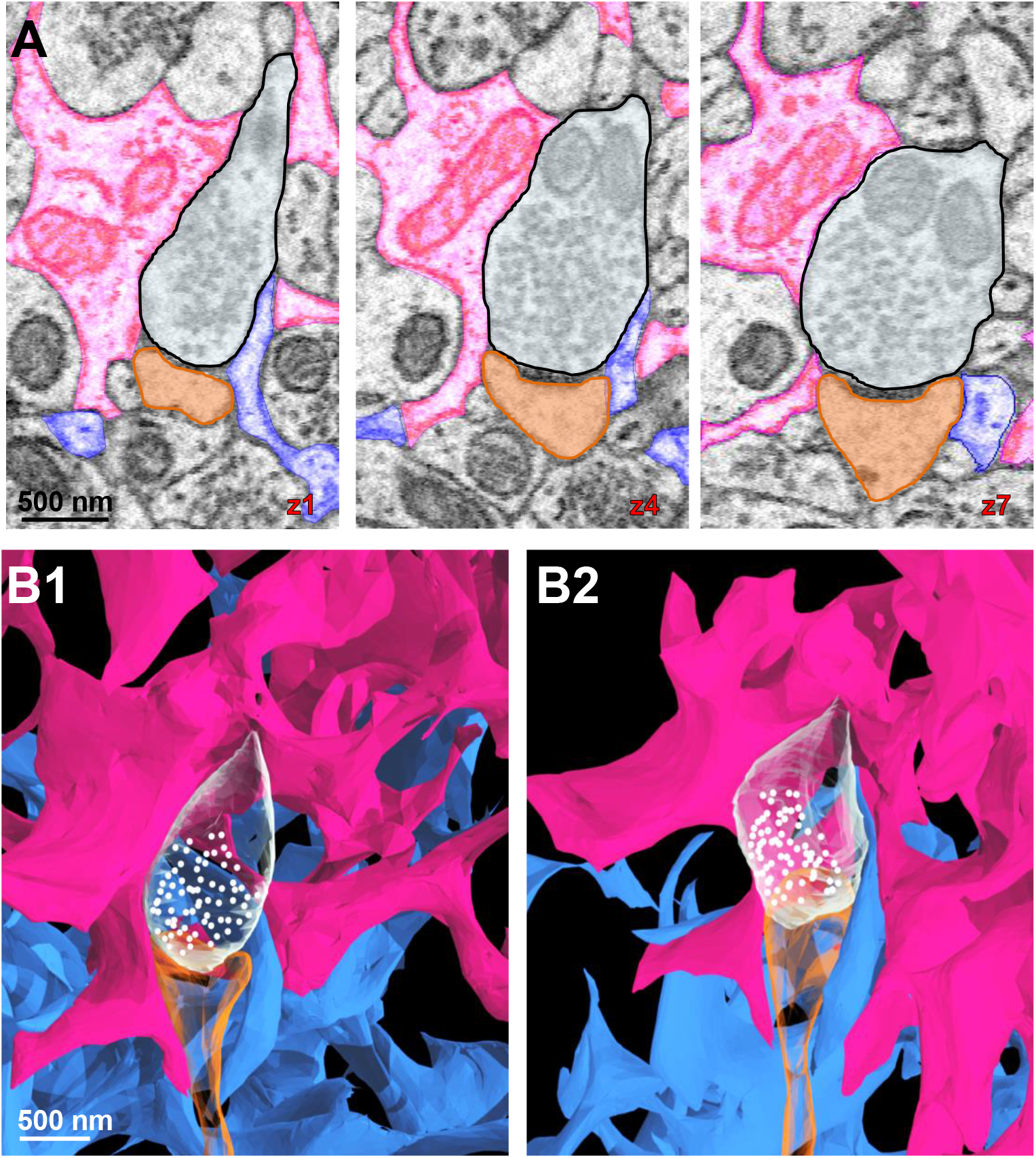
Synapses can be ensheathed by processes stemming from different astrocyte domains. **A)** 2D serial EM traces of two astrocyte processes (pink and blue) contacting the same synapse (white: axon; orange: dendritic spine head). **B1-B2)** 3D reconstructions of the two astrocytes contacting the same synapse (from ‘A’). Note that as shown in the 2D traces, the pink process contacts the synapse on the left side and the blue process on the right side.

## Discussion

Connectomics is an emerging subfield in neuroscience that aims to map the vast network of interactions within the brain (for excellent reviews, see Alivisatos et al., 2012; Fornito et al., 2015; Swanson and Lichtman, 2016). While the ultrastructure and connectivity of neurons has been highlighted in several seminal connectomics studies (Kasthuri et al., 2015; Mishchenko et al., 2010), our results provide significant insights into the structural complexity of astrocyte processes, the astrocyte-astrocyte contact patterns that underpin a low electrical resistance pathway, and the anatomical relationship between astrocytes and synapses in the mature mouse brain. Taking advantage of an *Aldh1l1*-eGFP mouse for pre-identification of neighboring astrocytes in EM specimen preparation and SBF-SEM for preserving the nanoscopic details of astrocytic processes, the first ultrastructural view of an astrocyte connectome is provided in this study.

### Complexity of astrocyte processes: spongiform morphology and neurite structural support

With respect to astrocyte process patterning, they generally followed a Root-Intermediate-Terminal (RIT) scheme, which closely resembles dendritic topology that has been previously reported (Uylings and van Pelt, 2002). However, two distinct patterning features were observed. First, numerous processes (with size and width comparable to that of terminal processes) extended out from every part of astrocyte (Fig. 2C). Second, we observed an abundance of reflexive, loop-like structures. Although technical limitations (in resolution) precluded us from validating that these self-connecting structures contain gap junctions, other studies have confirmed the existence of gap junctions in such reflexive contacts (Nagy et al., 1997; Wolff et al., 1998). These two features together give rise to spongiform morphology of astrocytes in our connectome (Video S1).

The significant number of astrocyte reflexive processes raises interesting questions about their functional contribution. Notably, we observed these processes ensheath both dendritic and axonal structures (Fig. 2 G-H), signifying their structural support role for neurites. In addition, these loop-like structures increase the surface area so that astrocyte processes have a higher capacity to interact with more synapses. This view is supported by our observation that reflexive astrocyte processes accommodated the second most synapses (with ~0.95 synapses per unit area) compared to all other astrocyte process types (Fig. 5). This unique ability for astrocytes to loop around neurites may provide structural fluidity, such that astrocytes can accommodate and support existing synapses. Along these lines, a recent study revealed that astrocyte ‘nodes’ (which often formed similar ‘loop-like’ structures) are the sites of initiation of calcium signals. Indeed, calcium activity at the nodes positively correlated with dendritic spine size, suggesting that these structures make up the astrocytic component of the tripartite synapse (Arizono et al., 2020). Furthermore, the structure of these astrocytic reflexive processes is similar to that of mesaxons - extensions of the Schwann cell membrane (Peters, 1964, 1960) which contain gap-junction-like structures (Bertaud, 1978; Mugnaini et al., 1977; Sandri et al., 1977; Tetzlaff, 1982) and allow for the transport of ions and other small particles across the myelin sheath (Balice-Gordon et al., 1998). The structural similarity between our observed astrocytic reflexive processes and Schwann cell mesaxons raises the prospect that both process types exert comparable functional roles.

### Anatomical and biophysical basis of low inter-astrocytic pathway resistance

The idea that low-resistance pathways exist between astrocytes was first discovered over 50 years ago in mudpuppy optic nerves (Kuffler et al., 1966). Recently, a gap junction mediated low inter-astrocyte electrical resistance was directly demonstrated in rodents and that serves as the *biophysical* basis for a phenomenon termed syncytial isopotentiality (Ma et al., 2016). However, until now, the *anatomical* basis underlying the low resistance pathway between astrocytes was not known.

Our data confirmed that astrocyte-astrocyte contacts occur only at their terminal processes, or the interface of astrocytic domains (Bushong et al., 2002; Ogata and Kosaka, 2002). Further, the ultrastructural details of two major inter-astrocytic contact types, point-point and elongate, were revealed. Analysis of these contacts allowed us to biophysically rationalize how anatomical inter-astrocyte contact patterning leads to low resistance. Notably, in our mathematical modeling of a full ‘connectome unit’ (with 6 different contacts; Fig. 3D), the coupling coefficient (CC) was simulated to be 88.17%, which is below the experimentally tested CC of 94% (Ma et al., 2016). Hence, assuming that this mix of contacts is the standard ‘connectome unit’ occurring in the astrocyte-astrocyte interface, our computational modeling predicts that three connectome units are sufficient to achieve this experimentally validated 94% CC.

We should also note that the abundance of reflexive loops and the structural variability of these loops made quantification difficult; hence, our mathematical simulations were made without the consideration of reflexive terminal processes in order to maintain confidence in our modeling. Biophysically, the reflexive loops should be another mechanism by which the astrocyte is able to lower its overall pathway resistance. This can be achieved through an increase in the transverse area of the totality of the astrocyte process, effectively enlarging the diameter of the cable. Hence, fewer ‘connectome units’ would be required in order for reflexive terminal processes to achieve the experimentally validated 94% CC.

While our current study did not elucidate the functional significance of the various types of astrocyte contacts, the assembly of astrocytes into a low-resistance syncytium raises the question as to whether this anatomical design endows astrocytes the ability to more closely associate with synapses in order to modulate brain function. Along these lines, the conductivity of gap junctions can be regulated through various signaling pathways as a result of neurotransmitter influx and pH or temperature change (for reviews, see Bukauskas and Verselis, 2004; Goodenough and Paul, 2009). For example, in cultured astrocytes, glutamate was found to potentiate gap junction coupling (Enkvist and McCarthy, 1994), suggesting that the coupling strength of a ‘connectome unit’ can be regulated by synaptic activity. How the conductivity of gap junctions could be modulated by other neurotransmitters is not fully understood; nevertheless, these findings raise the prospect that neurotransmission can enhance or weaken coupling within a given connectome unit. Hence, at any given time, certain connectome units within an interface may undergo strengthening while other units may weaken. As a result, syncytial isopotentiality is a spatiotemporal summation of all events occurring across all connectome units, and it is likely influenced by the proximity of astrocytes and synapses (a topic discussed below).

### Ultrastructural contacts of astrocytes with synapses

By tracing all axons from three dendrites, we were able to analyze astrocyte-synapse association, reporting - for the first time within the adult mouse hippocampus - that ~86% of synapses are ensheathed by astrocytes. Notably, this percentage is higher than other studies that examined synapse coverage within the developing rat somatosensory cortex (68% coverage) (Kikuchi et al., 2020) and the mature rat stratum radiatum (57% and 62% coverage, respectively) (Ventura and Harris, 1999; Witcher et al., 2007). Differences in brain region, developmental stage, and/or analysis methods likely contributed to the increase in astrocyte-ensheathed synapses that we observed. Further, our finding that a large majority of astrocytes contact the axon-spine interface (i.e. synapse), compared to post or pre-synaptic elements, is in agreement with these previous studies in both the developing (Kikuchi et al., 2020) and mature brain (Witcher et al., 2007). Of note, whereas most asymmetric synapses are excitatory, symmetric synapses are inhibitory (Peters and Palay, 1996). Interestingly, ~94% of all synapses reported in the P14 developing brain (Kikuchi et al., 2020) were asymmetric, a finding that we also observed in our study in the mature (P45) brain, suggesting that the instructive role of astrocytes in synaptogenesis occurs mainly in the developing brain (Allen and Eroglu, 2017).

In terms of astrocyte-synapse interaction, a current view is that astrocytes extend their terminal processes (PAPs) to ensheathing synapses; these PAPs provide structural support, clear neurotransmitters from the synaptic cleft, and respond to synaptic activity (Araque et al., 2014; Papouin et al., 2017). Hence, PAP is considered as the third anatomical and functional component in synaptic transmission, termed ‘tripartite synapse’ (Araque et al., 1999). Interestingly, while we noted that the density of synapses surrounding terminal processes was very high, we found that that all other astrocytic cellular compartments (soma, root, intermediate and reflexive processes) were also able to contact synapses. These observations indicate that every subcellular structure an astrocyte is equally equipped with the same synaptic support machinery as terminal processes, and thus, the same functions assigned to the canonical ‘PAPs’ may also apply to the entire astrocyte.

In our 3D reconstructions of network-level astrocyte-synapse contacts, we observed synapses being ensheathed by more than one astrocyte. These observations provide clear anatomical evidence that synaptic coverage does not occur exclusively within one astrocyte domain - a finding that was also observed using multi-color electron microscopy (Adams et al., 2016). These results also suggest that astrocytes within a given region are unlikely to be functionally specialized. Likewise, because synapses can associate with more than one astrocyte, they too, could exert various functional roles depending on the support they receive from each astrocyte.

### Paucity of vesicle-like structures within astrocytes

The fact that astrocytes contact synapses is central to the highly debated topic of gliotransmission (Fiacco and McCarthy, 2018; Savtchouk and Volterra, 2018), wherein astrocytes modulate synaptic transmission via calcium-dependent vesicular release of ‘gliotransmitters’ (i.e. neurotransmitters, ATP, neuropeptides). Hence, if one were to postulate that astrocytes do, in fact, transfer information to neurons through a vesicular release mechanism, then we would expect to observe concentrated clusters of vesicles in astrocytes adjacent synapses - similar to the abundance of vesicles observed in the presynaptic axon terminal. In contrast, however, our analysis revealed that only a few vesicles were located in astrocytes and that they do not specifically abut synapses. Furthermore, we show that synapse contact with astrocytes is ubiquitous, as synapses make contact with the astrocyte soma, root, intermediate, and terminal processes. Meanwhile, we observed a lack of vesicular-like structures throughout the entirety of the astrocyte (Fig. 7), which is in agreement with a recently published study reporting that astrocyte processes do not contain structures similar to neurotransmitter vesicles (Chai et al., 2017). Furthermore, the existence of a quantal release system - similar to that observed in synapses (Augustine and Neher, 1992; KATZ, 1959; Pankratov et al., 2007) - remains controversial in astrocytes (Pangrsic et al., 2007; Pascual et al., 2005; Stout et al., 2002; Xiong et al., 2018b; Zhang et al., 2007)(for review, see Xiong et al., 2018a). Thus, while it is unlikely that gliotransmission operates through astrocyte vesicular exocytosis, several EM studies have reported L-glutamate, D-serine, and VGLUT3 labeled synaptic-like microvesicles within astrocyte compartments (Bergersen et al., 2012; Ormel et al., 2012). Although the quantity of these vesicle-like structures observed within astrocytes in our study (and in others) is small, this is only a problem if it is assumed that the transmitter release sites of astrocytes must be the same as those found in nerve terminals – which is an unlikely scenario given that astrocytes are not excitable and cannot, therefore, activate any reserve pool like neurons (Savtchouk and Volterra, 2018). Hence, caution should be taken when considering this paucity of astrocyte vesicles as clear evidence against gliotransmission. Clearly, our understanding of the extent to which astrocytes are able to transfer information to neurons is still in its infancy.

## Acknowledgements

This work was sponsored by grants from National Institute of Neurological Disorders and Stroke; Grant code: RO1NS062784, R56NS097972, RO1NS116059 (MZ), and National Science Foundation; Grant Code: DMS1410935 (DT).

## Author Contributions

CMK, SA, GK, DT, MZ designed the experiments, CMK, SA, EPA, ATT, YD, AMG, MP, EGC, DM, KC, EB, and JAP conducted the experiments; CMK, SA, EPA, ATT, JAP analyzed the data; SA, CMK, MZ wrote the paper; all authors revised and approved the final draft of the paper.

## Tables

**Supplementary Table 1 (Table S1).** Synaptic density in relation to astrocyte process type. Tukey post-hoc test was conducted to determine statistically different synapse densities across all astrocyte process types. This data supports standard ANOVA analysis from Fig. 5H.

| **Synaptic Density-Astrocyte Process Comparison** | **Adjusted P- Value** | **Significance** |
| --- | --- | --- |
| Soma vs. Root process | 0.1840 | ns |
| Soma vs. Intermediate process | 0.0009 | *** |
| Soma vs. Terminal process | <0.0001 | **** |
| Soma vs. Reflexive process | <0.0001 | **** |
| Soma vs. Endfeet process | 0.9111 | ns |
| Root process vs. Intermediate process | 0.3187 | ns |
| Root process vs. Terminal process | <0.0001 | **** |
| Root process vs. Reflexive process | 0.0231 | * |
| Root process vs. Endfeet process | 0.8522 | ns |
| Intermediate process vs. Terminal process | 0.0017 | ** |
| Intermediate process vs. Reflexive process | 0.7967 | ns |
| Intermediate process vs. Endfeet process | 0.0457 | * |
| Terminal process vs. Reflexive process | 0.0756 | ns |
| Terminal process vs. Endfeet process | <0.0001 | **** |
| Reflexive process vs. Endfeet process | 0.0024 | ** |

**Supplementary Table 2 (Table S2).**
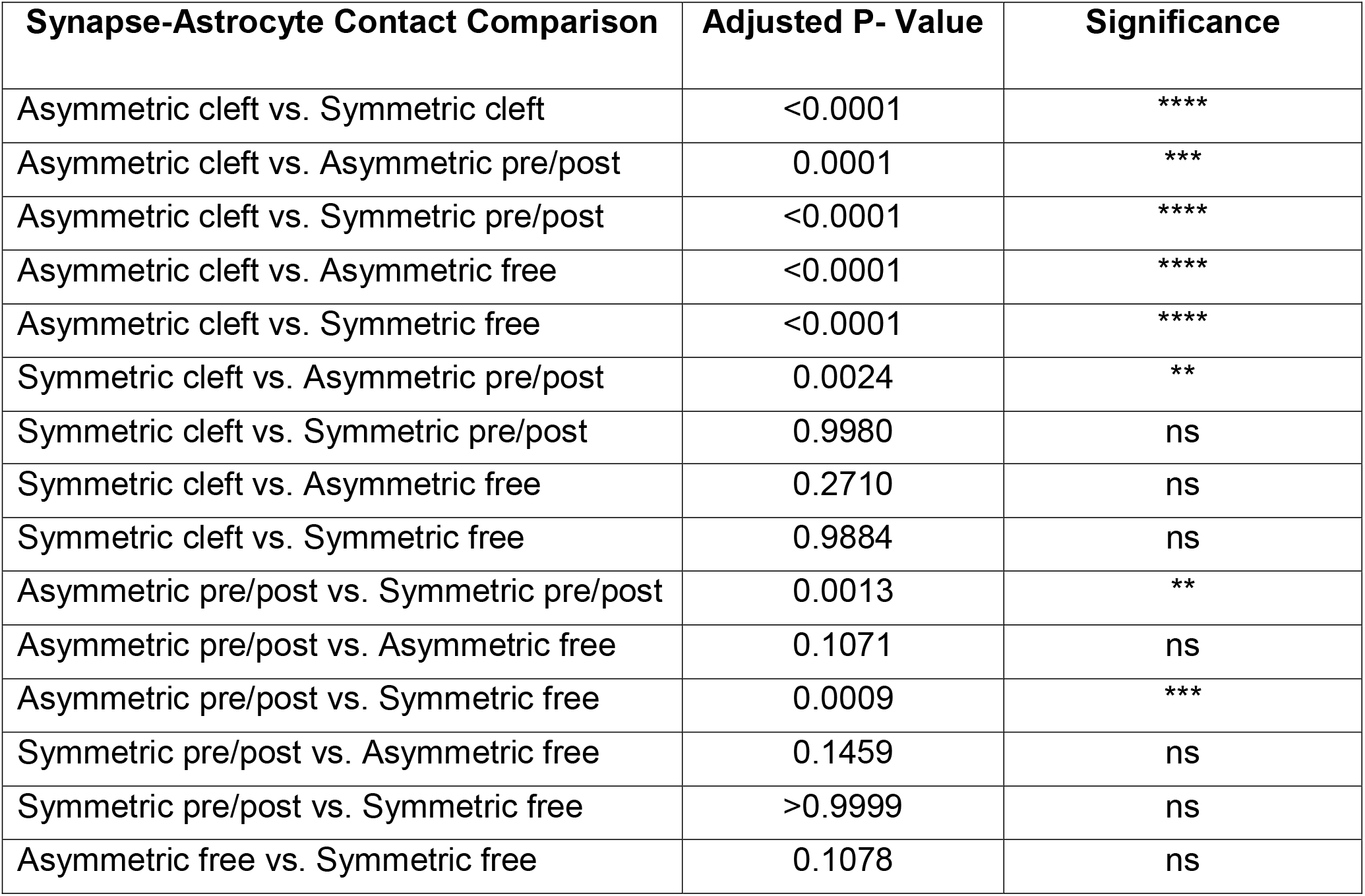
Asymmetric and symmetric synapse-astrocyte contact information from all 920 analyzed synapses. Tukey post-hoc test was conducted to determine statistically different synapse-astrocyte contacts (as a function of synapse type: i.e. asymmetric vs. symmetric). This data supports standard ANOVA analysis from Fig. 6E1.

| Synapse-Astrocyte Contact Comparison | Adjusted P- Value | Significance |
| --- | --- | --- |
| Asymmetric cleft vs. Symmetric cleft | <0.0001 | **** |
| Asymmetric cleft vs. Asymmetric pre/post | 0.0001 | *** |
| Asymmetric cleft vs. Symmetric pre/post | <0.0001 | **** |
| Asymmetric cleft vs. Asymmetric free | <0.0001 | **** |
| Asymmetric cleft vs. Symmetric free | <0.0001 | **** |
| Symmetric cleft vs. Asymmetric pre/post | 0.0024 | ** |
| Symmetric cleft vs. Symmetric pre/post | 0.9980 | ns |
| Symmetric cleft vs. Asymmetric free | 0.2710 | ns |
| Symmetric cleft vs. Symmetric free | 0.9884 | ns |
| Asymmetric pre/post vs. Symmetric pre/post | 0.0013 | ** |
| Asymmetric pre/post vs. Asymmetric free | 0.1071 | ns |
| Asymmetric pre/post vs. Symmetric free | 0.0009 | *** |
| Symmetric pre/post vs. Asymmetric free | 0.1459 | ns |
| Symmetric pre/post vs. Symmetric free | >0.9999 | ns |
| Asymmetric free vs. Symmetric free | 0.1078 | ns |

## STAR Methods

### Key Resources Table

Key Resources Table:
| REAGENT or RESOURCE | SOURCE | IDENTIFIER |
| --- | --- | --- |
| **Deposited Data** |  |  |
| Raw and analyzed data | This paper | N/A |
| **Experimental Models: Organisms/Strains** |  |  |
| BAC *Aldh1l1*-eGFP mice | GENSAT Project at Rockefeller University | MGI:3843271<br>IMPC: Tg(Aldh1l1-EGFP)OFC789Gsat |
| **Software and Algorithms** |  |  |
| Reconstruct | Synapse Web | RRID: SCR_002716 |
| Blender 2.78 and 2.90 | https://www.blender.org/download/releases/2-78/ | RRID:SCR_008606 |
| GraphPad Prism 7 | https://www.graphpad.com/scientific-software/prism/ | RRID:SCR_002798 |

### Supplementary Videos

**Supplementary Video 1 (Video S1). 3D video reconstruction of 3 neighboring astrocytes.**

3-dimensional, rotation view of a blood vessel (red) and three reconstructed astrocytes: purple, blue, and pink. This video supports reconstructions observed in Fig. 1I-K.

**Supplementary Video 2 (Video S2). 3D video reconstruction of a reflexive astrocyte process enwrapping a dendrite.**

3-dimensional view of a reflexive astrocyte process (pink) ensheathing a dendritic stalk (orange). Also note the reflexive contact depicted in yellow. This video supports reconstructions shown in Fig. 2G2.

**Supplementary Video 3 (Video S3). 3D video reconstruction of a reflexive astrocyte process enwrapping axons.**

3-dimensional view of a reflexive astrocyte process (purple) ensheathing a bundle of axons (white). Also note the reflexive contact depicted in yellow. This video supports reconstructions shown in Fig. 2H2.

**Supplementary Video 4 (Video S4). 3D video reconstruction of inter-astrocyte contact patterns.**

3-dimensional view of a ‘point-point’ contact and an ‘elongate’ contact between two astrocytes (blue and pink), in addition to a full ‘connectome unit.’ This video supports reconstructions and computational modeling depicted in Fig. 3B2, C2, and D.

**Supplementary Video 5 (Video S5). 3D video reconstruction of 3 neurites.**

3-dimensional, rotation view of three dendrites (orange) and the axons (white) that contact each dendritic spine head. This video supports reconstructions observed in Fig. 4A-C.

**Supplementary Video 6 (Video S6). 3D video reconstruction of 3 astrocytes and 3 neurites.**

3-dimensional, rotation view of a blood vessel (red), 3 dendrites (orange) and associated axons (white), and the three reconstructed astrocytes (pink, purple, and blue). This video supports reconstructions observed in Fig. 4D.

**Supplementary Video 7 (Video S7). 3D video reconstruction of two astrocytes supporting the same synapse.**

3-dimensional view of two astrocytes (pink and blue) ensheathing the same synapse (orange: post-synaptic element; white: pre-synaptic element). This video supports reconstructions shown in Fig. 8B1-B2.

### Lead Contact

Further information and requests for resources and reagents should be directed to and will be fulfilled by the Lead Contact, Dr. Min Zhou.

### Materials Availability

This study did not generate any new/unique reagents.

### Data and Code Availability

The full EM Reconstruct files (i.e. tracings) will be deposited in a public repository upon acceptance of the manuscript.

### Experimental Model and Subject Details

An adult, postnatal day 45, female BAC *Aldh1l1*-eGFP mouse was used in this study. Details of this mouse line have previously been reported (Yang et al., 2011). Mice were housed in a temperature controlled (22 ± 2°C) environment with a 12 hour light/dark schedule and ad libitum access to food and water. All procedures were performed in accordance with a protocol approved by the Institutional Animal Care and Use Committee (IACUC) at The Ohio State University.

### Method Details

#### Tissue processing

A postnatal day (P) 45 mouse was anesthetized with an intraperitoneal injection of 8% chloral hydrate in 0.9% saline and then transcardially perfused at 6 mL/min with 4% paraformaldehyde and 2.5% glutaraldehyde in 0.1M sodium cacodylate buffer. Sodium cacodylate, paraformaldehyde, and shell vials were purchased from Electron Microscopy Sciences (Hatfield, PA, USA). 25% EM grade glutaraldehyde was purchased from Polysciences Inc (Warminster, PA, USA). Coronal CA1 hippocampal brain slices (300 μm) were then cut with a Vibratome (Pelco 1500) and post-fixed with the same fixative overnight (at 4 degrees C) in shell vials.

#### Correlative confocal and serial blockface scanning electron microscopy (SBF-SEM)

Each *Aldh1l1*-eGFP hippocampal brain slice was then imaged using a Leica (SP8) confocal microscope in order to obtain astrocyte spatial localization and blood vessel landmark visualization within the stratum radiatum. These confocal images were used to correlate and select a region of interest for subsequent electron microscopy tissue processing.

After confocal images were acquired, fixed tissue sections were washed five times in 0.1 M sodium cacodylate, followed by staining with reduced osmium (2% osmium tetroxide and potassium ferrocyanide in 0.1 M sodium cacodylate buffer) at 4C for 2.5 hours. Sections were then washed five times in double distilled water, followed by incubation in 1% thiocarbohydrazide at 60C for 1 hr, before being washed again in double distilled water. Sections were then stained with 2% aqueous osmium tetroxide for 2 hr at room temperature and were subsequently incubated (for 24-30 hr) in aqueous uranyl acetate at 4C. Next, the tissue was washed and incubated for 60 minutes at 60C in Walton’s lead aspartate, followed by washing in double distilled water before beginning dehydration through a series of incubations in ethyl alcohol, propylene oxide (4 hrs), and 90 min in Epon 812-substiute resin before being embedded in Epon 812-substitute resin and left to cure for 48 hr at 60C. Next, the tissues were trimmed out of the resin and oriented on the pin according to the corner notch that was cut into the wet tissue before confocal imaging. The tissue was mounted on aluminum and coated with colloidal silver liquid around the exposed edges of the resin block. A Zeiss Sigma VP system (with an in-chamber Gatan 3 View ultamicrotome with low-kV backscattered electron detectors) was used to examine the tissue. Tissue samples were imaged at 2.2 kV, with 7.7 nm/pixel resolution. Slices were 75 μm thick. SBF-SEM image acquisition and registration was conducted at Renovo Neural (Cleveland, OH, USA). The total image scan size for the data set is 54.02 × 96.47 × 37.5 μm (X, Y, Z). Image series were registered and then analyzed using Reconstruct (see below).

#### Three-dimensional reconstructions of hippocampal astrocytes, neurites, blood vessels, and intracellular particles

500 serial SBF-SEM images of the stratum radiatum (provided by Renovo Neural) were imported with image pixel size (0.0077 μm/pixel) and slice thickness (0.075 μm) into Reconstruct (Fiala, 2005). Cellular structures were traced manually or traced using the Reconstruct wildfire tool in order to generate individual objects. Structures traced with the wildfire tool were checked and manually corrected. The volume of each completed object was automatically generated in Reconstruct. Further, the Z-trace tool was used to measure the dimensions (size, length, etc.) of each object, and the ellipse tool was used to trace rounded objects (such as glycogen granules). Completed Boissonnat surface object reconstructions were generated in Reconstruct and were then exported as VRML 2.0 files for further rendering in Blender (see below).

#### Visualizing SBF-SEM reconstructed data using Blender 2.78

Blender - a free, open source 3D, general-purpose graphics tool that allows for modelling of large-volume data sets - was used to reconstruct our astrocyte connectome files. Note that Blender has been used in several other studies examining EM neural tissues (Calì et al., 2016; Zheng et al., 2018). VRML 2.0 files (created in Reconstruct) were imported into Blender and were then colored and rendered to obtain final 3D reconstructions. Note that in certain circumstances, objects were made slightly transparent (using the ‘Z transparency’ tool) to allow for easy visualization of multiple objects in contact with one another.

#### Blender video reconstructions and animations

Blender (v2.90.0) - an open-source software package - was utilized to model, render, and animate the rasterized 3-dimensional astrocyte-neurite network visualizations. The visualizations convey complex structures, and therefore specific rendering and staging techniques were used in the production process to ensure structures were visually distinct and to afford clarity to the represented forms. Along these lines, a two-source light design scheme was used for the videos, with one key light source positioned near the camera and a second source located in the subject’s rear. The key light source illuminates the subject’s front while reducing shadows that could obscure complex structures. The rear light source (referred to as a ‘hair light’) borrows from film lighting technique to accentuate and define very fine structures.

Further, care was taken to ensure that shadows did not obscure the complex structures in the reconstructed video animations. Toward this end, the lighting that was utilized simulates area lights which allow for volumetric shadows that mimic naturally occurring soft shadows and approximate the light source’s environmental diffusion, while also providing visual depth cues.

The rendered subject surfaces in the videos were achieved with two physically based rendering techniques - transmission and subsurface light scattering approximations - in order to achieve a volumetric design. These tactics act to soften the subject’s surface features and convey a waxy, organic appearance. Additionally, this strategy allowed for light transmission through the objects’ edges in order to create visual definition.

#### Mathematical modeling of astrocyte contacts

The Z-trace tool (Reconstruct) was used to measure the dimensions of astrocytic processes to be used in mathematical modeling simulations. To computationally model the effect of a cluster of astrocyte contacts on the coupling coefficient (CC), the area between the contacting astrocytes was measured using Reconstruct (these regions were the presumed sites of gap junctions). Further, the number of gap junctions was estimated based on the density of gap junctions from freeze-fracture replica immunogold labeling experiments (Rash et al., 2001).

The computational model was developed and simulated using the numerical software NEURON (Carnevale and Hines, 2006). The two astrocytes shown in Fig. 3D, bottom panel, consist of 12 (blue astrocyte) and 9 (pink astrocyte) compartments, respectively. Of these, there are single soma compartments, 2 and 1 root/primary process compartments, 2 and 1 intermediate/secondary process compartments, and 7 and 6 terminal/tertiary process compartments, respectively.

#### Synapse-astrocyte contact analysis

Analysis of astrocyte coverage of synapses was adapted from a recently published paper (Kikuchi et al., 2020). Note, however, that our analysis was conducted in SBF-SEM traces (i.e. 2-dimensional) and not in rendered (i.e. 3-dimensional) constructions. To begin, three complete dendrites (and their associated spines) were traced using Reconstruct. Axons that contacted each spine were also traced to completion. Next, each synapse was marked and classified as asymmetric or symmetric based on whether the postsynaptic density was prominent (Gray, 1959; Kikuchi et al., 2020; Peters and Palay, 1996) (see Fig. 6D1 for examples of the two types of synapses). Finally, the area surrounding each synapse was examined in order to determine whether any astrocyte processes were located adjacent to the synapse. Three categories were then established, similar to classifications conducted by Kikuchi et al (Kikuchi et al., 2020):
 
1. ‘Cleft associated astrocytes’ were those synapses whose synaptic clefts made contact with an astrocyte process.
2. ‘Pre/post associated astrocytes’ were those synapses whose pre or post synaptic elements made contact with an astrocyte process, but not with the cleft.
3. ‘Free astrocytes’ were those synapses that had no adjacent contacts with astrocyte processes.

Of note, if an astrocyte made contact with both the synaptic cleft and a pre or post synaptic element, the contact was classified as ‘cleft contact’. The percentage of astrocyte (and non-astrocyte) associated synapses was then calculated from three dendrites.

#### Analysis of vesicle-like structures within astrocyte processes

A 5 × 5 μm, 10 z-stack ROI was drawn around an astrocyte region (from a 2D EM trace). These regions included the astrocyte soma, root, intermediate processes, terminal and reflexive processes, and endfeet. Three of these ROIs were drawn for each astrocyte region. The vesicle-like structures within each ROI were counted, and the total number of vesicles within each ROI was divided by the total volume that the astrocyte process occupied in order to determine the density of vesicles per volume of astrocyte. This process was replicated for all process types of three different astrocytes (blue, pink, and purple). Criterion used to identify vesicle-like structures can be found in Fig. S1. As a control, the total number of synaptic vesicles was also analyzed using these same parameters - 5 × 5 μm boxes from synapses surrounding the astrocyte soma were marked as ROIs. The number of vesicles (in the stacks where the synapse was prominent) was divided by the volume of the presynaptic element (which contained the vesicles) in order to determine the density of vesicles.

#### Analysis of synapse contact with various astrocyte process types

Similar to the vesicle analysis (above), a 5 × 5 μm, 10 z-stack ROI was drawn around an astrocyte region (from a 2D EM trace). These regions included the astrocyte soma, root, intermediate processes, terminal and reflexive processes, and endfeet. Three of these ROIs were drawn for each astrocyte region. The synapses that contacted a region of the astrocyte within that ROI were traced and the total number of synapses that contacted the astrocyte within the 10 z-stack ROI was divided by the total volume that the astrocyte occupied. This process was repeated for all regions of three different astrocytes.

#### Statistical Analysis

All statistics were performed using GraphPad Prism 7.0 software, and group data are presented as mean ± SEM. Details of statistical tests (such as the test statistic, degrees of freedom, and p-value) can be found in the Results section (or supplementary tables) of the manuscript. As noted in the figure legends, significance was ascribed to p-values < 0.05. Further, comparisons between three (or more) groups/variables were conducted using a one-way ANOVA, followed by post-hoc analysis. For each experiment, Grubb’s test was conducted on data obtained from each group in order to determine whether the outlier was statistically significant (p < 0.05). Hence, this test was used to exclude one data point (terminal process ROI) from the analysis of vesicle-like structures within astrocytes (Fig. 7G). Additionally, Grubb’s test was used to exclude two data points (one soma ROI and one reflexive ROI) from the analysis of synapse density in relation to astrocyte process type (Fig. 5H).

---

**Supplementary Figure 1.**
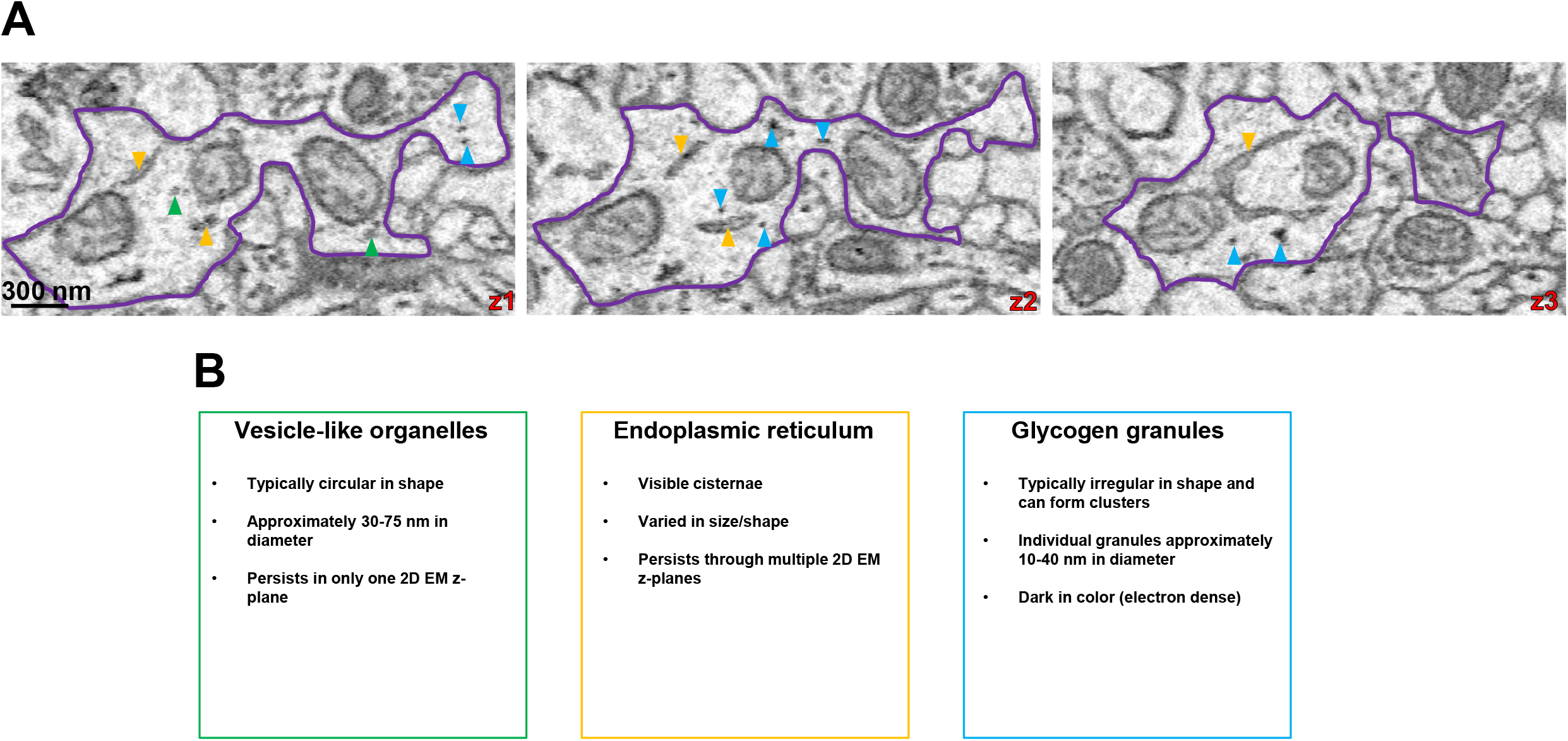
Criterion for identification of astrocytic intracellular particles. **A)** Serial 2D EM images of an astrocyte intermediate process containing vesicle-like particles (green arrowheads), endoplasmic reticulum (yellow/orange arrowheads), and glycogen granules/clusters (blue arrowheads). **B)** Classification of how each intracellular particle type was identified based on appearance, size, and morphology. Note that the color of the box corresponds to the color of the arrowhead in **A**.

**Supplementary Figure 2.**
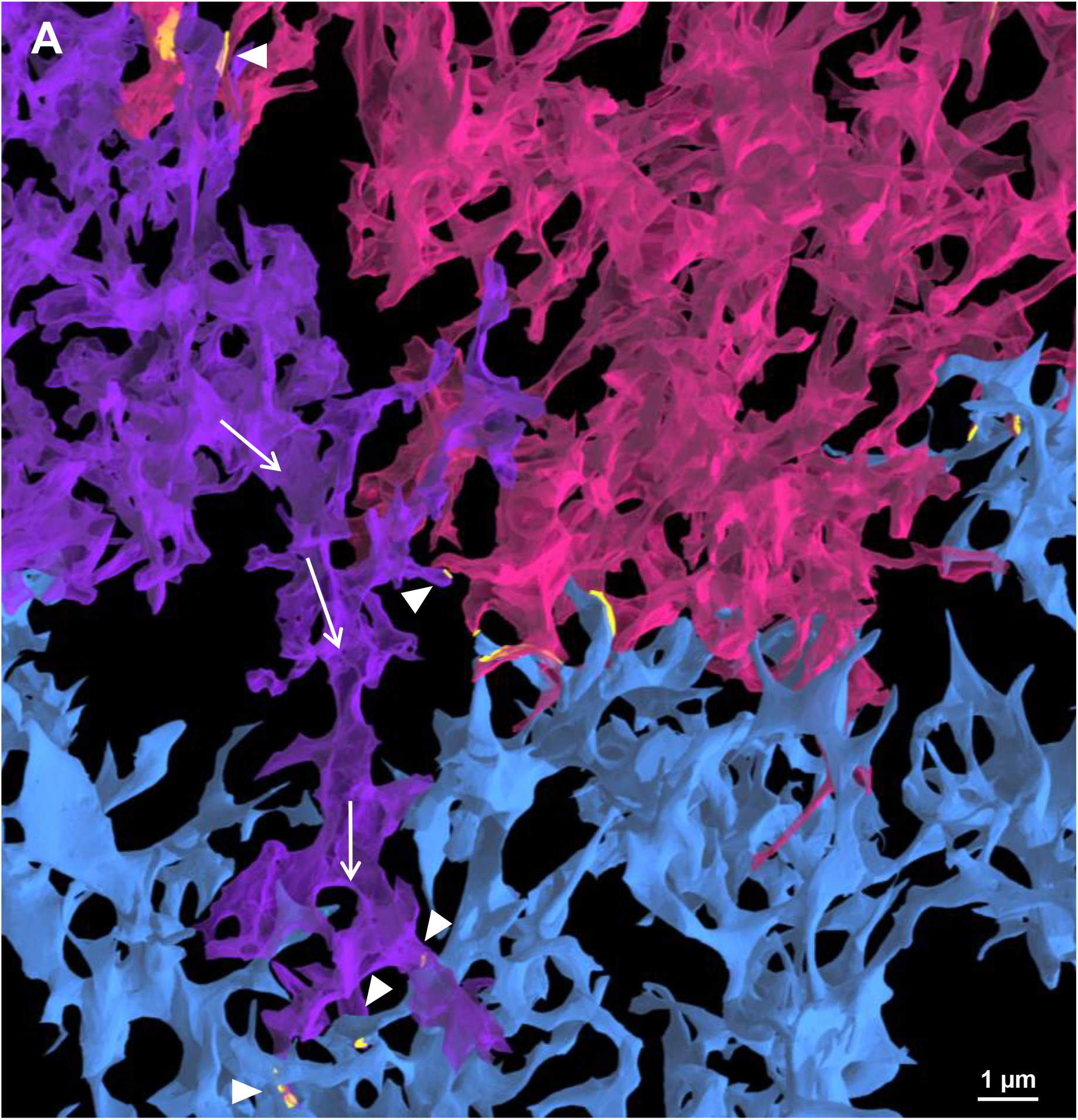
Processes from one astrocyte can contact more than one neighboring astrocyte. A) Low magnification view of the astrocyte connectome unit from Fig. 3A. Note that all three astrocytes (pink, blue, and purple) contact one another. The pink and purple astrocyte contact one another (top left; contact depicted in yellow), and as the purple astrocyte process extends downward (white arrows), it makes perpendicular contact with the pink astrocyte again (white arrowhead; contact in yellow), and finally extends to contact the blue astrocyte (white arrowhead; contact in yellow). The blue astrocyte also contacts both the purple and pink astrocytes, and the pink astrocyte contacts the blue and purple astrocytes.

**Supplementary Figure 3.**
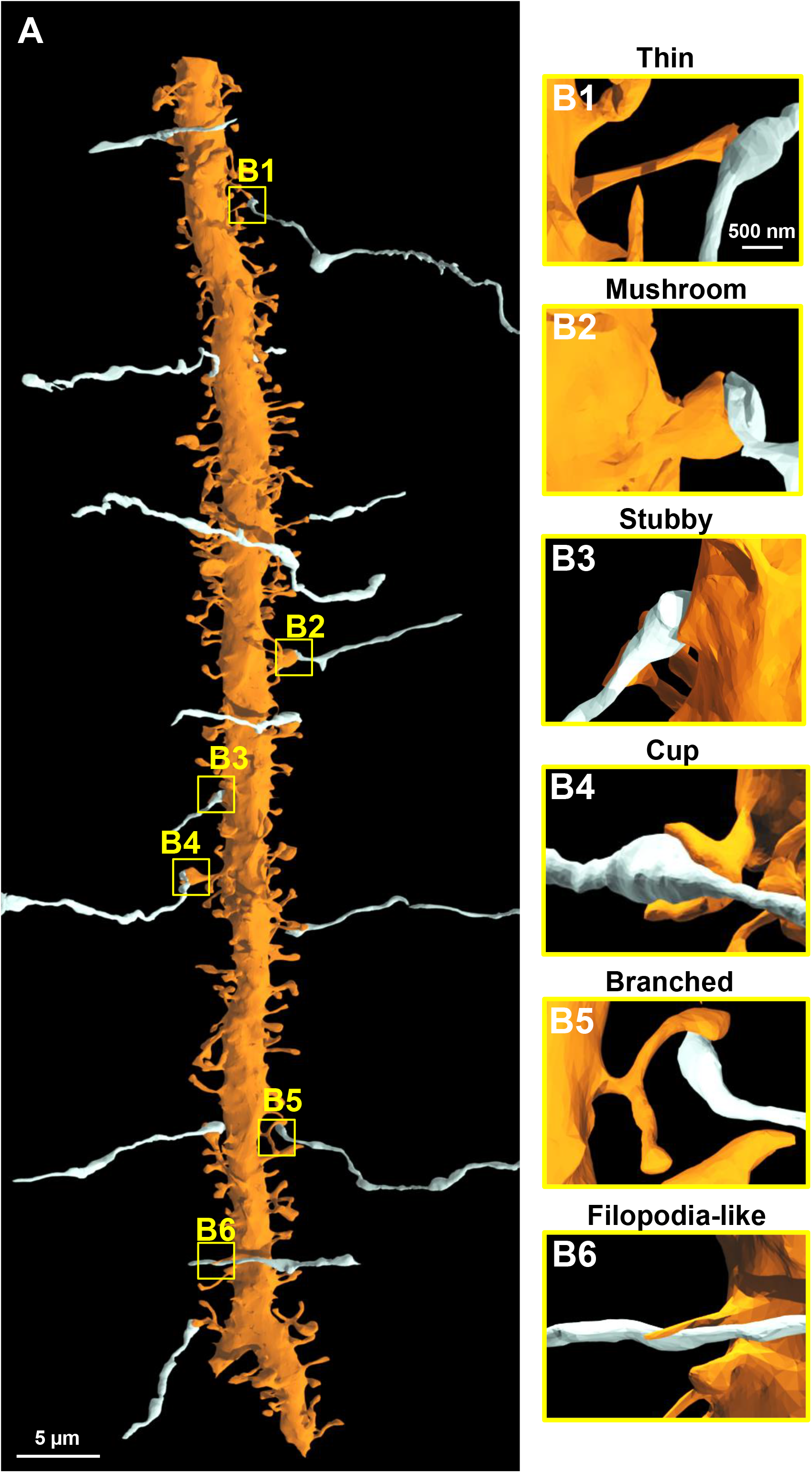
Ultrastructural view of a neurite. **A)** Complete 3D reconstruction of one (out of three total) dendrites. Several axons (white) are drawn for reference. **B1-B6)** Magnified view of axonal contacts (white) with dendritic spines (orange). The orange dendritic spines that contact these axons represent thin, mushroom, stubby, cup, branched, and filopodia-like spine types, respectively. Note that these images are magnified from yellow boxed regions approximated in **A**.

